# Evolutionary responses to historic drought across the range of scarlet monkeyflower

**DOI:** 10.1101/2025.05.20.655098

**Authors:** Seema Nayan Sheth, Lucas Albano, Charles Blanchard, Emily Cook, Rosalinda Diaz, Xitlaly Gomez-Vega, Katelin Kutella, Mariam Moazed, Macy Patel, Julia Prange, Niveditha Ramadoss, Ashley Regan, Aster Riley, Melissa Rivas Hernandez, Jocelyn Rojas, Marissa Strebler, Aditi Verma, Lindsay Villano, Jordan Waits, Dachuan Wang, Olivia Wilborn-Pilotte, Jeffrey Diez, Lluvia Flores-Rentería, Jason P. Sexton, Christopher D. Muir

## Abstract

Adaptive evolution is a key means for populations to persist under environmental change, yet whether populations across a species’ range can adapt quickly enough to keep pace with climate change remains unknown. The breeder’s equation predicts the evolutionary change in a trait from one generation to the next as the product of the selection differential and the narrow-sense heritability in that trait. Incorporating these aspects of the breeder’s equation, we performed a resurrection study with the scarlet monkeyflower (*Mimulus cardinalis*) to evaluate whether traits associated with drought adaptation have evolved in populations across a species’ range in response to extreme drought. We compared trait and fitness differences of pre-drought ancestors and post-drought descendants from six populations transplanted into three latitudinally-arrayed common gardens and quantified phenotypic selection and trait heritabilities. The strength, direction, and mode of selection varied among traits and gardens. Trait heritabilities were relatively low, and did not differ dramatically among populations or gardens. Overall, instances of evolutionary responses between ancestors and descendants were few and small in magnitude, but the magnitude of these evolutionary differences varied among gardens. Together, these results suggest that the expression of genetic variation, and thus traits, depend on the environment, and that environmental variability in field settings may mask the genetic variation that is often detected in greenhouse environments.

## Introduction

As human activities fragment landscapes and impede the movement of organisms, evolutionary adaptation may become increasingly necessary for populations to persist under environmental change. Ecologists and evolutionary biologists have long sought to understand how populations adapt to changing environments, rejuvenated by interest in predicting whether evolution can rescue populations from extinction due to climate change. Evolutionary rescue, the mechanism where a declining population becomes stable or growing due to adaptive evolution, directly bridges the evolutionary process of adaptation and the ecological process of demography (Gomulkiewicz & Holt 1995). The probability of evolutionary rescue depends on ecological and evolutionary factors (Bell & Gonzalez 2009; Carlson *et al*. 2014). For instance, larger population sizes, shorter generation times, and higher additive genetic variances for fitness-related traits increase the probability of evolutionary rescue for a population. Empirically, evolutionary rescue is often more likely to occur under gradual rather than abrupt environmental change in lab microcosms (Lindsey *et al*. 2013), but the likelihood of population persistence in nature is related to biological parameters, such as genetic variance, and the strength of phenotypic selection, which need to be measured in the field (Chevin *et al*. 2010). Since we lack good estimates of these parameters during extreme climatic events, we do not know whether adaptive evolution will rescue declining populations.

The amount of genetic variation that a population harbors for fitness-related traits should predict its rate of adaptation to environmental change and hence the probability of evolutionary rescue. The univariate (*R* = *h*^2^*S*) and multivariate (Δ*z* = **G**β) breeder’s equations predict the evolutionary change in one (*R*) or more (Δ*z*) traits from one generation to the next as the product of narrow-sense heritability (*h*^2^) and the selection differential (*S*) or the product of the additive genetic variance-covariance matrix (**G**) and the vector of selection gradients (β), respectively (Falconer & Mackay 1996; Lande & Arnold 1983). The breeder’s equation evaluates the rate of evolution for specific traits, but the potentially arbitrary choice about which traits to measure could lead to the omission of other fitness-related traits (Shaw 2019). Variation in the strength and direction of selection over time (Shaw 2019; Siepielski *et al*. 2009) and among different fitness components (Kingsolver *et al*. 2001; Siepielski *et al*. 2011) make it challenging in practice to predict phenotypic evolution in natural populations using the breeder’s equation.

Range limit theory suggests that the parameters involved in the breeder’s equation should vary predictably across species’ geographic ranges (reviewed in Sexton *et al*. 2009), leading to spatial differences in the probability of evolutionary rescue. When a species’ range is at equilibrium with its climatic niche (i.e., the range is stable and not contracting or expanding), theory predicts that fitness and standing genetic variation should decrease from the range center towards the edges (Pironon *et al*. 2016), and edge populations should experience stronger directional selection than central populations (Sexton *et al*. 2009). Populations can harbor substantial genetic variation for ecologically important traits (Etterson & Shaw 2001; Mousseau & Roff 1987), facilitating rapid adaptation (Bush *et al*. 2019; Hairston *et al*. 2005), but it is untested whether range limit theory accurately predicts adaptive potential among populations (Pennington *et al*. 2021). As ongoing climate change decouples range limits from climatic niche limits, trailing range edges at equatorial latitudes and downslope elevations may have low population growth rates and standing genetic variation. Conversely, leading range edges at poleward latitudes and upslope elevations could have higher population growth rates and standing genetic variation than trailing range edges due to gene flow from migrating populations (Davis & Shaw 2001; Hampe & Petit 2005; Pironon *et al*. 2016). Additional biological parameters such as genetic drift can further complicate these simplistic predictions (Polechová 2018, 2025). Further, since the magnitude and direction of climate anomalies can vary across space and time (Burrows *et al*. 2011), the strength, direction, and mode of selection can vary among populations across a species’ range (Munguía-Rosas *et al*. 2011; Siepielski *et al*. 2013, 2017). In addition to the spatial variation in *h*^2^ and *S* described above, populations across species’ ranges can vary in their evolutionary responses (*R*) to artificial selection on ecologically important traits (Kelly *et al*. 2012; Pujol & Pannell 2008; Sheth & Angert 2016). Thus, the probability of evolutionary rescue likely differs across species’ ranges. High population growth rates and standing variation in the range center and incoming variation to leading range edges suggest these may be hotspots for evolutionary rescue, while trailing edge populations, if suffering from continued erosion of genetic variation due to niche-limiting climate stress, may be coldspots for evolutionary rescue. Nevertheless, trailing edge populations may still be hotspots of unique genetic variation as isolated and relictual, post-glacial refugia (Hampe & Petit 2005). We need empirical studies that assess genetic variation, the strength of natural selection, and evolutionary responses to that selection in multiple range edges within species as predicted by range limit theory (Angert *et al*. 2020).

Many regions of the world are experiencing increased temperatures, intensified droughts, and greater temporal variability in temperature and precipitation (Diffenbaugh & Field 2013; Swain *et al*. 2018). In herbaceous plants, the organismal focus of this study, populations may adapt by timing their phenology to escape stressful conditions, or evolve to physiologically avoid stress in increasingly harsh and variable environments (Franks & Hoffmann 2012; Hoffmann & Sgro 2011; Kooyers 2015). In plant evolutionary ecology, escape refers to shifts in seasonal timing, whereas avoidance refers to physiological shifts in optimal performance (Franks 2011; Kooyers 2015; Ludlow 1989). We do not discuss stress tolerance (e.g., the ability to maintain turgor pressure under severe drought) (Aslam *et al*. 2015) because it is probably not common in herbaceous plants (Kooyers 2015). The likelihood of evolutionary response in escape or avoidance traits depends on the relative strength of selection and additive genetic variance for each trait (Bradshaw & Holzapfel 2008; Hoffmann & Sgro 2011). For example, *Brassica rapa* populations evolved earlier flowering times in response to a period of drought, and plants that flowered earlier had lower water-use efficiencies, indicative of tradeoffs between the evolution of escape and avoidance traits (Franks 2011). In the absence of tradeoffs, if one only measured escape traits, but avoidance traits responded to selection, this would lead to the wrong conclusion of no evolutionary response. Ultimately, we need to understand evolutionary responses in both escape and avoidance traits to determine if herbaceous plant populations can persist *in situ*.

Here, we evaluate the degree to which drought escape and avoidance traits have evolved in populations across a species’ range in response to extreme climatic events. To do so, we take advantage of a well-studied, broadly distributed species (*Mimulus cardinalis*, syn. *Erythranthe cardinalis*) in which populations vary significantly in climatically-relevant phenotypes, genetic variance in these phenotypes, vital rates and their contributions to population growth rate, and the magnitude of environmental change (Anstett *et al*. 2024; Branch *et al*. 2024; Muir *et al*. 2022; Muir & Angert 2017; Nelson *et al*. 2021; Sheth & Angert 2016, 2018). We performed a resurrection study comparing ancestors and descendants derived from seed collected before and after a 7-year period of severe drought and heat. We transplanted pre-drought ancestors and post-drought descendants from populations from across the species’ range into three common gardens at contrasting range positions and quantified traits associated with drought escape and avoidance along with first-year fitness. Adapting the breeder’s equation, we estimated *h^2^* of each trait and phenotypic selection on each trait in each garden to assess their potential influences on the evolution of trait differences between ancestors and descendants. We also addressed the following hypotheses (Fig. 1). First, extreme drought caused evolution in escape or avoidance traits, but the magnitude differs between leading- and trailing-edge populations based on range-limit theory (H1). Second, the magnitude of evolutionary response differs across space (i.e., among gardens) because the expression of genetic variation is environmentally-dependent (Hoffmann & Merilä 1999; H2). For instance, populations may exhibit reduced evolutionary responses under warmer, drier conditions if stress increases environmental variation (Bemmels & Anderson 2019; Charmantier & Garant 2005; Hoffmann & Merilä 1999). This work demonstrates how combining a resurrection study with common gardens can provide robust examinations of predictions of evolutionary change based on quantitative genetic theory (Wadgymar *et al*. 2023).

**Figure 1.**
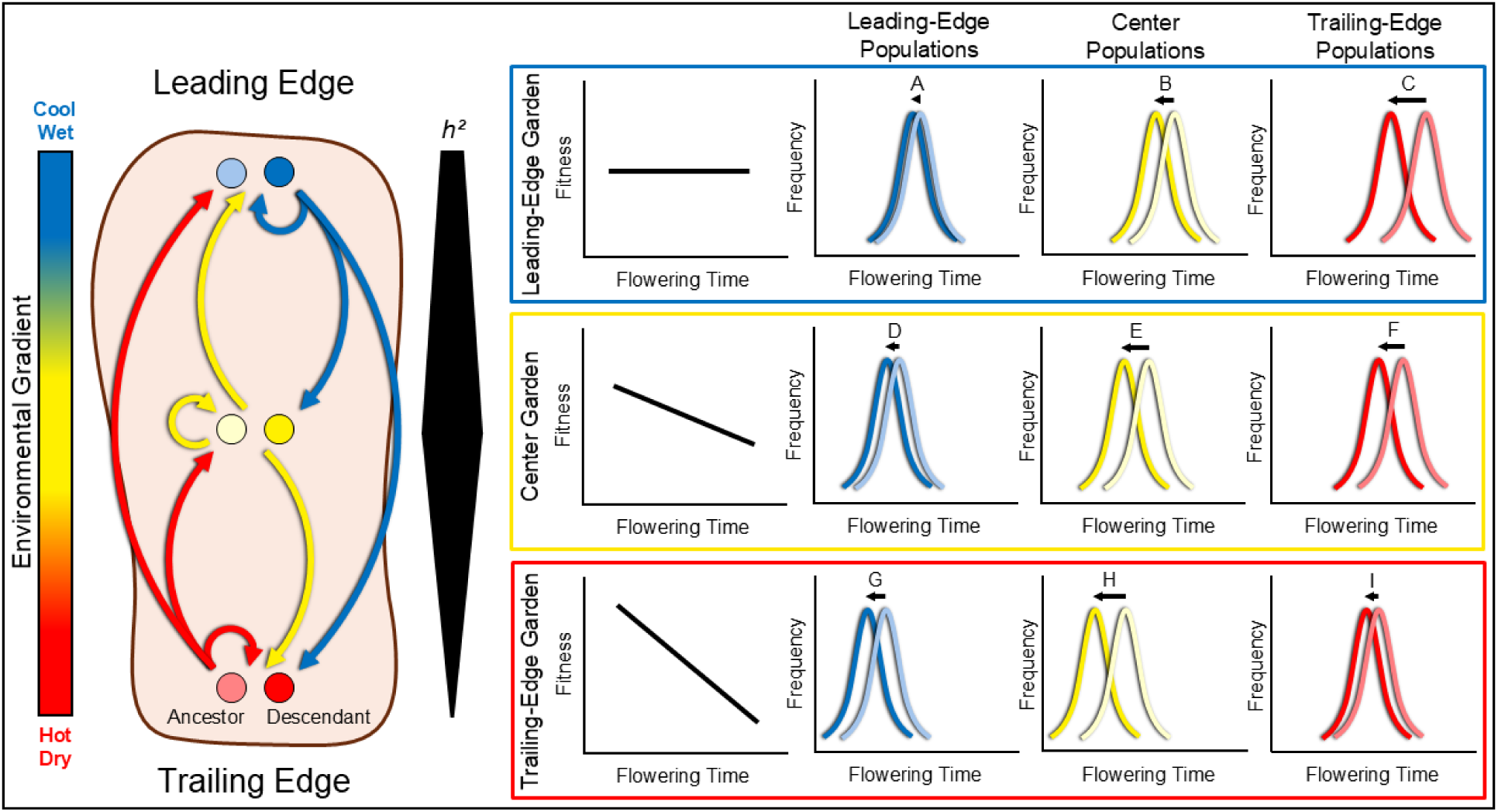
Illustration of the hypotheses addressed by combining resurrection and space-for-time approaches (Franks *et al*. 2014; Karitter *et al*. 2024; Lovell *et al*. 2023) in experimental field gardens spanning an environmental gradient. Ancestors (lighter hues) and descendants (darker hues), originating from propagules collected before and after an episode of environmental change, from populations at the leading edge (blue), center (yellow), and trailing edge (red) of a species’ range are transplanted into common garden sites (represented by the colored rectangles surrounding the graphs) in these same three areas of the range. Range limit theory yields alternative predictions about how genetic variation (and thus heritability, *h^2^*) and the strength of directional selection (quantified via the selection differential, *S*) vary across species’ ranges (Sexton *et al*. 2009). This example illustrates one possible scenario. In this scenario, *h^2^* is highest in the range center and lowest at the trailing edge (identified by the width of the black bar labelled *h²*), and the strength of directional selection (based on *S*) declines from the trailing edge towards the leading edge. We illustrate selection on flowering time, one ecologically important trait associated with adaptation to the environmental gradient, as an example. Given this scenario, the magnitude of evolutionary change in flowering time between ancestors and descendants should vary among populations originating from across the species’ range (e.g., A vs. B vs. C; H1). Leading-edge, central, and trailing-edge gardens exhibit variation in the magnitude of evolutionary response between ancestors and descendants due to environmentally-dependent expression of genetic variation among transplanted populations (e.g., A vs. D vs. G; H2). Further, based on the breeder’s equation, (*R* = *h²S*), the relative magnitude of evolutionary response between ancestors and descendants of each population within gardens would be expected to be proportional to standing genetic variation (e.g., G vs. H vs. I) and the strength of selection (e.g., A vs. B vs. C). In this example, if spatial patterns of phenotypic selection across the range predict temporal evolutionary responses between ancestors and descendants associated with climate change, then leading-edge and central populations should exhibit evolution of advanced flowering from ancestors to descendants when transplanted into warmer, drier environments (e.g., D, G, and H).

## Methods

### Study system

We used the scarlet monkeyflower (*Mimulus cardinalis*, Phrymaceae) as a model system for studying evolutionary responses to climate change. *Mimulus cardinalis* is a perennial herb that grows in riparian areas from central Oregon, USA, to northern Baja California, Mexico (Fraga, 2018; Lowry et al., 2019), spanning a broad climatic and latitudinal gradient (Fig. 2).

**Figure 2.**
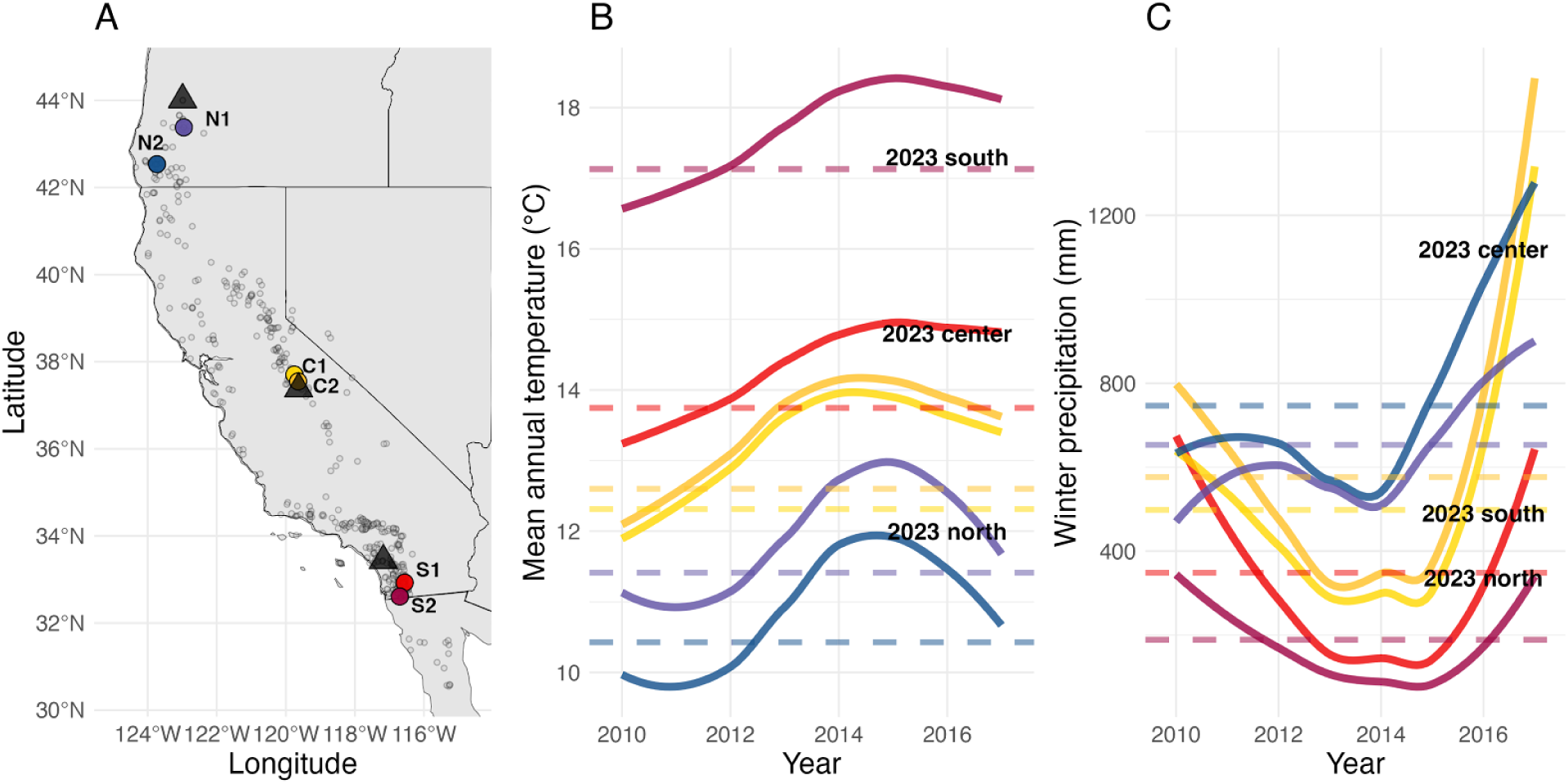
A) Map of known *Mimulus cardinalis* occurrences (open circles, from (Angert *et al*. 2018)), focal populations (filled circles), and common garden locations (black triangles). B) Mean annual temperature for each focal population from 2010 to 2017 (curves), along with historical 30-year averages (horizontal dashed lines, 1981 - 2010). C) Winter precipitation for each focal population from 2010 to 2017 (curves), along with historical 30-year averages (horizontal dashed lines, 1981 - 2010). Climate data for each focal population are derived from climateNA v. 7.30 and data for each garden are derived from the web version of climateNA v. 7.60 (Wang *et al*. 2016). In panels B and C, the text denotes mean annual temperature (B) and winter precipitation (C) for each garden in 2023.

*Mimulus cardinalis* is a classic evolutionary ecology system (Hiesey *et al*. 1971; Schemske & Bradshaw 1999) for investigating geographic range limits (Angert 2006a, b, 2009; Angert *et al*. 2008, 2018; Angert & Schemske 2005; Bayly & Angert 2019; Paul *et al*. 2011) and potential physiological (Angert *et al*. 2011; Branch *et al*. 2024; Muir & Angert 2017), evolutionary (Anstett *et al*. 2021; Sheth & Angert 2016; Vtipil & Sheth 2020; Wooliver *et al*. 2020), and demographic (Anstett *et al*. 2024; Sheth & Angert 2018) responses to climate change. From 2012 to 2015, western North America experienced multiple years of record warming compounded by severe drought (Diffenbaugh *et al*. 2015; Robeson 2015; Wang *et al*. 2016), with variation in the magnitude of drought among populations across the range of *M. cardinalis* (Fig. 2b, c).

Range-wide demographic surveys overlapping with this historic drought (2010-2014) found multiple years in which high mortality and/or no seedling recruitment occurred, leading to significant declines in 19 out of 32 populations spanning the latitudinal gradient (Sheth & Angert 2018). The populations experiencing the most anomalously dry conditions in winter showed the greatest declines during the drought (Anstett *et al*. 2024).

The evolutionary response to climate change likely differs across the range of *M. cardinalis* due to strong growth and survival responses to temperature and drought (Angert 2006b; Muir & Angert 2017; Paul *et al*. 2011; Wooliver *et al*. 2020) and latitudinal variation in escape and avoidance traits and demography (Muir & Angert 2017; Nelson *et al*. 2020; Sheth & Angert 2016, 2018). Previous work reveals the potential for rapid evolutionary responses in escape and avoidance traits in *M. cardinalis*. For instance, broad-sense heritabilities in some life history and leaf functional traits are substantial (Muir *et al*. 2022; Nelson *et al*. 2020). A greenhouse experiment including the ancestral cohort of the same focal populations as the present study suggests potentially adaptive variation in flowering time, measured as the number of days from germination to first flower, across the species’ range, whereby flowering time increased from north to south (Sheth & Angert 2016). When we artificially selected for early and late flowering time, southern populations responded rapidly, whereas northern and central populations showed negligible responses. Thus, there is potential for inter-population variability in evolutionary response to severe drought and warming, with southern populations having ample genetic variation for flowering time to adapt, whereas the response in northern and central populations could be limited. Selection for early flowering resulted in correlated increases in specific leaf area and leaf nitrogen content, consistent with a drought escape strategy (Sheth & Angert 2016). Conversely, selection for late flowering resulted in correlated responses in drought avoidance traits. Thus, climate change may lead to shifts along the escape-avoidance continuum in this species.

### Focal populations

We focus on two leading-edge (northern), two central, and two trailing-edge (southern) populations across the geographic range of *M. cardinalis* (Fig. 2a; Sheth & Angert 2016; Wooliver *et al*. 2020). We collected seeds from ∼50-200 individuals from each population in 2010 (ancestral cohort) and 2017 (descendant cohort). In May 2018, we propagated field-collected seeds from both cohorts of the 6 populations for a “refresher generation” to control for maternal and storage effects (following Franks *et al*. 2007) in a greenhouse at North Carolina State University. At least 80% of families and ∼68-92% of total field-collected seeds planted from each population-cohort combination germinated (Vtipil & Sheth 2020), consistent with high seed survival and minimal invisible fraction bias (Franks *et al*. 2019; Weis 2018). We performed crosses within each population and cohort to produce seeds of known pedigree with a nested paternal half-sibling design (8-33 sires per population-cohort combination x 3-5 dams/sire = 1266 full-sib families nested within 268 half-sib families across all populations and cohorts). This design allows phenotypic variation to be partitioned among sires, among dams nested within sires, and within dams. The number of sires varies among populations due to differences in population size across the species’ range–we later standardized total sample sizes among populations (see below).

### Experimental design

To test the hypotheses above, we used the broad geographic range of *M. cardinalis* to perform a space-for-time substitution (Franks *et al*. 2014; Karitter *et al*. 2024; Lovell *et al*. 2023). Northern-edge populations have already begun to encounter temperatures and levels of drought comparable to those experienced historically by southern-edge populations (Sheth & Angert 2018). We established common gardens in three positions within the range of *M. cardinalis*: near the northern range limit (Friends of Buford Park and Mt. Pisgah Native Plant Nursery; Eugene, OR), the latitudinal center (Sierra National Forest’s Batterson Work Center; Oakhurst, CA), and the southern range limit (San Diego State University’s (SDSU) Santa Margarita Ecological Reserve; Temecula, CA; Fig. 2a).

We used the pedigreed seeds from each population and cohort to establish seedlings in greenhouses at UO, UCM, and SDSU in spring of 2023. To ensure that quantitative genetic parameters for all population-cohort combinations were based on similar sample sizes, we used the same number of individuals per group (population × cohort), and adjusted the number of replicates per full-sib family depending on the number of available full-sib families per group. Because large sample sizes are required for estimating quantitative parameters in the ancestral cohorts (Conner & Hartl 2004; Falconer & Mackay 1996), sample sizes for ancestors are greater than those of descendants. Specifically, we included 10 replicates per full-sib family for populations with < 25 sires, and 6 replicates per full-sib family for populations with ≥ 25 sires for each ancestral cohort, and 200 seedlings per descendant cohort per population per garden. To achieve a sample size of 200 seedlings for each descendant cohort, we randomly selected a maximum of 25 sires from each population with > 25 sires, and accordingly adjusted the number of dams and the number of replicates per dam, and for populations with < 25 sires, we included all full-sib families and replicated each family 2-3 times. Because of limited space in the central garden, we only planted sufficient full- and half-sibling replication to estimate quantitative genetic parameters for the ancestral cohorts of the two central populations. All other populations and cohorts in the central garden had similar sample sizes and sire and dam inclusion as the descendant cohorts in the other gardens. Plant trays were sub-irrigated and misted daily in the greenhouse during the first week to promote seedling establishment, and then sub-irrigated daily until transplanting in the field.

When the seedlings were ∼4-6 weeks old, we transplanted them using a randomized block design such that each sire from each population and cohort was represented in each block in each garden (14,290 total plants representing an average of 21 sires and 101 dams per ancestral cohort of each population; Tables S1, S2). Individual plants were spaced ∼0.3m apart in 4 rows per block (except one block in the central garden had 3 rows), and each block had a ∼0.5m-wide walkway on either side. In the southern garden, we installed below-ground gopher fencing, and all gardens were surrounded by deer fencing. In all gardens, we laid out landscape fabric and burned planting holes in the fabric with a blowtorch. We used micro-perforated drip tape (Aqua-Traxx, Toro, El Cajon, CA, USA) to irrigate each row of plants in each garden. We irrigated 5-7 days per week initially to promote the establishment of seedlings, which typically grow in wet riparian areas. As the growing season progressed, we gradually reduced irrigation and terminated irrigation upon the onset of winter precipitation. Additional details about each garden are in Table S1.

The climate during our study in 2023 did not reflect drought conditions, since our study occurred during a year with historic winter precipitation that occurred before we transplanted seedlings (Table S1, Fig. 2b, c). Notably, the central garden was wetter than the northern and southern gardens, and the northern garden in Oregon actually received less winter precipitation than the southern garden in 2023 (Table S1, Fig. 2b, c). Yet, climatic moisture deficit, a measure of drought severity that integrates temperature and precipitation, was highest in the central and southern gardens and lowest in the northern garden in 2023 (Table S1). Since 2023 was a climatically anomalous year that did not reflect longer-term climatic variation across the range of *M. cardinalis,* we do not focus on hypotheses that rely heavily on space-for-time substitutions in this study. We transplanted seedlings in the central garden ∼1 month later than in the southern garden and ∼2 months later than in the northern garden (Table S1) due to high snowfall and the associated delays in winter and spring site access and development.

### Data collection

Throughout the 2023 growing season, we measured several drought escape and avoidance traits (Kooyers 2015). In general, drought escape is associated with fast growth and development in order for plants to reproduce before the onset of severe heat and drought. “Escape” phenotypes therefore include earlier initiation of flowering at a smaller plant size, resource-acquisitive (high specific leaf area) and less heat/drought tolerant (low leaf dry matter content) leaves, and stomatal conductance that is high early in the growing season and low late in the growing season. Conversely, drought avoidance is associated with slower development and hardier growth in order for plants to withstand the effects of severe heat and drought later in the growing season. “Avoidance” phenotypes therefore include later initiation of flowering at a larger plant size, resource-conservative (low specific leaf area) and more heat/drought tolerant (high leaf dry matter content) leaves, and stomatal conductance that is low early in the growing season and high late in the growing season.

To assess the timing of flowering and size at first flower, we conducted semi-weekly phenology surveys throughout the growing season in each garden. Each plant was assessed for the presence of open flowers and the length (cm) of the three longest stems at the time of flowering (summed to obtain a more comprehensive estimate of size at first flower). We also assessed each plant for mortality and excessive damage, such as nearing mortality with some live tissue still present, or breakage of primary stems. To measure leaf traits and early-season (north: May 23 - May 31; south: June 6 - June 16) stomatal conductance, we selected the upper-most, fully expanded, healthy, and undamaged leaf on a main stem of each plant. Since plants in the central garden went into the ground somewhat later than the already delayed natural phenology in the relatively wet growing season for this region in 2023, we did not collect data on early-season physiological or leaf traits in the central garden. Stomatal conductance (*g_sw_*; in mmol/m²/s) was measured adjacent to the midvein at the distal end of the abaxial side of each selected leaf using a porometer (LI-600, LI-COR Environmental, Lincoln, NE, USA). Leaves were then harvested and scanned with a flatbed scanner to obtain leaf area (in cm²) measurements using FIJI version 2.14.0 (Schindelin *et al*. 2012). Fresh mass was measured in grams using an analytical balance (Model SPX123 Scout Balance, Ohaus, Parsippany, NJ).

Leaves were then dried at ∼76°C for at least 48 hours and weighed again. We calculated specific leaf area (*SLA*, cm²/g) as: *leaf area / leaf dry mass*, and leaf dry matter content (*LDMC*, g/g) as: *leaf dry mass / leaf fresh mass*. To assess late-season (north: August 26 - August 30; center: August 22 - August 24; south: August 15 - August 18) stomatal conductance, we again followed the leaf selection and stomatal conductance protocol above, but at all three gardens. Plants that were too small, unhealthy, or had no available undamaged leaves were excluded from collection of leaf trait and stomatal conductance data. At the end of the growing season, we recorded the height of the longest stem (in cm) and estimated the total number of reproductive structures (buds, flowers, fruits, and pedicels) on each plant as measures of size and fitness. We used the number of reproductive stems and the number of reproductive structures on up to three stems to estimate the total number of reproductive structures on each plant (calculations and validations in Supplementary Methods and Results, Table S3, Figs. S1, S2). Some rows/blocks were also excluded from some aspects of data collection due to time constraints (Table S4).

### Statistical analyses

For all models described below, we used the “brm” function in the R v. 4.5.0 (R Core Team 2025) package “brms” version 2.22.0 (Bürkner 2017, 2018), which allows Bayesian statistical analysis using Hamiltonian Monte Carlo in Stan version 2.35.0 (Carpenter *et al*. 2017) via the “cmdstanr” version 0.8.1 backend (Gabry *et al*. 2024). The “brms” package implements a wide variety of statistical distributions for non-Gaussian traits (e.g. binomial for survivorship, negative binomial for fecundity) and statistically accounts for common issues in field data such as zero-inflation and missing observations. To ensure parameter convergence, determined from a modified Gelman-Rubin statistic 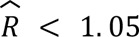 (Vehtari *et al*. 2021), we ran each model with four parallel chains for 1000 warmup iterations and 1000 sampling iterations per chain with no thinning. We used default priors in “brms”: improper flat prior for fixed effects, except the model intercept which assumed a Student-*t*(3, 0.5, 2.5) prior; weakly informative half Student-*t*(3, 0.5, 2.5) priors for random effect standard deviations; weakly informative LKJ(1) priors for random effect covariance matrices. Most parameters converged (Rhat < 1.05, ESS > 1000), but some constrained parameters and random effect coefficients did not quite meet these convergence criteria. Constrained parameters (e.g. variances must be greater than 0 and correlation coefficients are bounded by −1 and 1) often do not meet these convergence criteria without more informative priors, but we prefer to use weakly informative priors despite suboptimal performance. The few random effect coefficients that failed to converge are minor and do not meaningfully impact our main inferences. All parameter estimates and confidence intervals were summarized using the median and 95% quantile intervals of the posterior distribution, respectively.

#### Analyses of trait differences between ancestors and descendants of each population in each garden (H1 and H2)

To explore patterns of trait variation and test for evolutionary responses between ancestors and descendants of each population in each garden, we built Gaussian linear mixed effects models of each trait as a function of population, garden, cohort and all possible interactions as fixed effects. We included sire nested within each population, garden, and cohort, dam nested within sire, and garden block as random effects in each model. We also modeled residual standard deviation separately for each population and garden to obtain population-specific narrow-sense heritability estimates in each garden (see below). For the early- and late-season stomatal conductance models, we also included log-transformed leaf vapor pressure deficit and time of day of data collection (with a quadratic term due to non-linearity in these variables and scaled and centered to improve numerical stability and computational efficiency) as fixed effects, and date of data collection as a random effect because stomatal conductance can vary with vapor pressure deficit, time of day, and date. In these models with additional fixed effects, we also included an interaction between time of day and garden, and all possible interactions between log-transformed vapor pressure deficit with the other fixed effects. For the model for total number of reproductive structures, we added a constant of 1 (to avoid log_10_(0)) and then log_10_-transformed this trait. We used 95% confidence intervals around trait medians to evaluate how each trait varied among populations and gardens. To assess evolutionary responses of each trait in each population in each garden (H1 and H2), we determined whether the 95% confidence intervals for each trait difference between descendants and ancestors overlapped 0.

#### Estimation of *h^2^*

The total phenotypic variation (*V_P_*) of the ancestral cohort of each population in each garden can be partitioned into genetic and environmental variances (*V_G_* and *V_E_*, respectively). *V_G_* can be further partitioned into additive genetic variance (*V_A_*) and non-additive genetic variance, which includes both dominance and epistatic variance (*V_D_* and *V_I_*, respectively). A nested paternal half-sibling design allows for the estimation of the observational variance components of sire 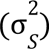, dam 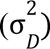, and error (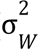; within-progeny) variances, which sum to the total phenotypic variance (Conner & Hartl 2004; Falconer & Mackay 1996). We used the models described above to estimate *V_A_* in each trait for the ancestral cohort of each population in each garden as 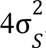. We estimated environmental variance (*V_E_*) for each population in each garden from the residual standard deviation, and then estimated narrow-sense heritability (*h^2^*) as the ratio of *V_A_* to *V_P_* (the sum of sire, dam, and error variance). To determine whether there was significant *V_A_* in each population, we used a leave-one-out cross-validation information criterion (*LOOIC*) calculated with the “loo_compare” function in the “brms” package (Vehtari *et al*. 2017) to compare the full model above to a model without sire and dam random effects. To assess whether populations and gardens differed in *V_A_*, we compared the full model to models of separate *V_A_* for each population and garden, respectively. As a robustness check, we also compared these estimates of *h^2^*from global brms models to estimates derived from sub-models (e.g., the ancestral cohort of a single population in a single garden) in both brms and the more commonly used package, MCMCglmm (Hadfield 2010) (Supplementary Methods and Results, Table S5, Fig. S3). To visualize potential genotype-by-environment interactions (*G x E*) in greater depth, we plot breeding values of each source population across each pair of gardens, focusing on flowering time as an example (Sheth *et al*. 2018) (Supplementary Methods and Results, Fig. S4).

#### Selection analyses

To determine whether the strength, direction, and mode of selection on each trait varied among gardens, we modeled the log_10_-transformed total number of reproductive structures (+1 to avoid log_10_(0)) as a function of each trait (with a quadratic term to test for stabilizing selection), population, garden, and cohort as fixed effects, and the random effects of sire nested within each population, garden, and cohort, dam nested within sire, and garden block. We also modeled residual standard deviation separately for each population and garden to mirror the model structure that we used for the models of each trait above. We included an interaction between the trait and garden to account for possible differences in the strength and form of selection among gardens, and all possible interactions between garden, population, and cohort. In each univariate selection model, the trait was scaled and centered to improve numerical stability and efficiency, and we log-transformed late-season stomatal conductance prior to scaling and centering. We doubled quadratic regression coefficients to obtain stabilizing selection gradients (Stinchcombe *et al*. 2008).

## Results

### Variation in traits among populations and gardens

On average, early-season physiological and leaf traits associated with drought avoidance were more prevalent in the northern garden, and escape traits were more common in the southern garden. Specifically, plants in the northern garden overall had higher *LDMC* and lower *SLA* and early-season *g*_sw_ relative to plants in the southern garden (Table S6, Fig. 3). Conversely, phenological, late-season physiological, and fitness traits were not consistent with this pattern.

**Figure 3.**
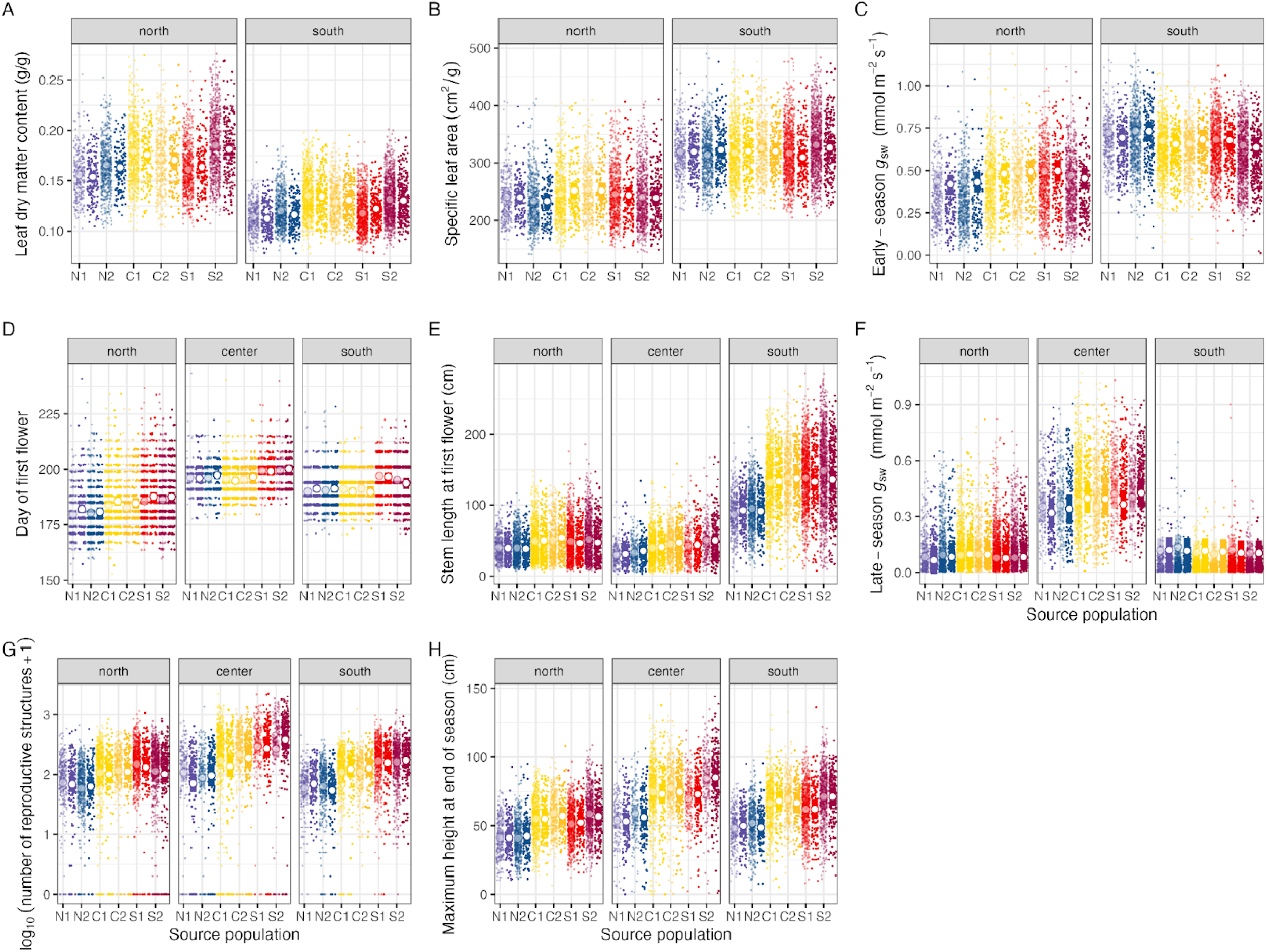
Model-predicted traits of ancestors (lighter hues) and descendants (darker hues) of each source population from the northern range edge (N1, N2), latitudinal range center (C1, C2), and southern range edge (S1, S2) in each common garden. White circles represent median values, and bars correspond to 95% confidence intervals. Colored points represent observed data. Stomatal conductance: *g_sw_*. Logistical constraints prevented us from collecting data on leaf traits and early-season *g_sw_* in the central garden.

Flowering time was earliest in the northern garden, intermediate in the southern garden, and latest in the central garden (Table S6, Fig. 3). Stem height at first flower was highest in the southern garden and lowest in the central garden (Table S6, Fig. 3). Late-season *g*_sw_, the total number of reproductive structures (our fitness proxy), and maximum height at the end of the growing season were highest in the central garden (Table S6, Fig. 3).

The degree of genetic differentiation among populations varied by trait and garden and sometimes opposed the direction of among-garden differences. For instance, though *LDMC* was lower in the southern versus the northern garden, it tended to increase from northern to southern populations within each garden (Table S6, Fig. 3). In the northern garden, flowering time was earliest for northern populations, intermediate for central populations, and latest for southern populations, while in the central and southern gardens, flowering time for northern and central populations was a similar magnitude earlier than southern ones (Table S6, Fig. 3). Stem height at first flower was greater in central and southern populations relative to northern populations in each garden (Table S6, Fig. 3). At the end of the growing season in each garden, the southern populations tended to make the most reproductive structures and have the tallest maximum height, whereas the northern populations made the fewest reproductive structures and were shortest (Table S6, Fig. 3). *SLA* and early- and late-season *g*_sw_ did not differ drastically among populations within gardens (Table S6, Fig. 3).

### Patterns of selection in each garden

The strength and mode of selection varied among traits and gardens, and selection did not consistently favor a single drought strategy in any garden. *LDMC* was not under selection in the northern garden, and was under weak stabilizing selection in the southern garden (Table S7, Fig. 4). *SLA* was not under selection in the north, and exhibited weak, negative directional selection in the south. (Table S7, Fig. 4). In the northern garden, early-season *g*_sw_ was under stabilizing selection, whereas there was positive directional selection in the southern garden. Selection favored early flowering in all gardens, and was strongest in the southern garden and weakest in the central garden (Table S7, Fig. 4). Size at first flower was under stabilizing selection in all gardens, with the most positive linear coefficient in the south, and the most negative quadratic coefficient in the center. Late-season *g*_sw_ was under stabilizing selection in the northern and central gardens, and positive directional selection in the southern garden (Table S7, Fig. 4). Our fitness proxy increased with maximum height at the end of the growing season in all three gardens, but began to decrease in the tallest plants, leading to stabilizing selection in all gardens, and the strength of directional selection was highest in the central garden and lowest in the northern garden.

**Figure 4.**
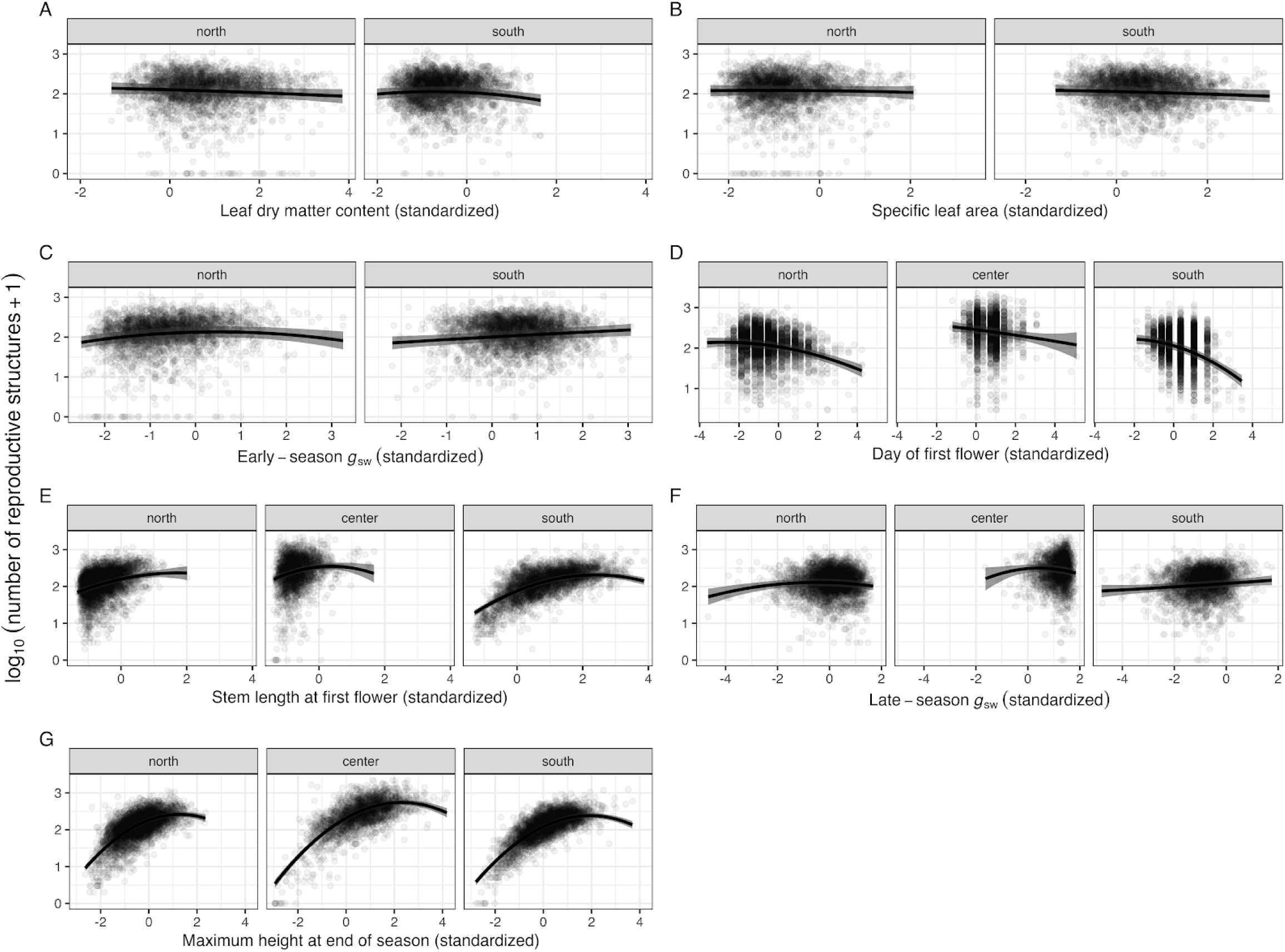
Log_10_-transformed fitness (measured as the number of reproductive structures) regressed on each trait in each garden. The black line is the slope of the regression between each trait and fitness, estimated from the median of the posterior distribution. The gray ribbon represents the 95% confidence intervals of the slope. Points represent individual plants. Each trait was scaled and centered prior to analyses, and late-season stomatal conductance (*g_sw_*) was log-transformed. Logistical constraints prevented us from collecting data on leaf traits and early-season *g_sw_* in the central garden.

### Variation in ancestral h^2^ among populations and gardens

Narrow-sense heritability in most traits was largely similar among populations and gardens and the lower 95% CIs were often close to 0 (Table S8, Fig. 5). In the northern garden, N2 had higher *h^2^* for *LDMC* than S2 (Table S8, Fig. 5). For all traits, the model with separate *V_A_* for each garden but no differences in *V_A_* among populations performed best, and for every trait except late-season *g*_sw_, the model without sire and dam effects performed the worst (Table S9), revealing substantial *V_A_* that varied among gardens, but not populations.

**Figure 5.**
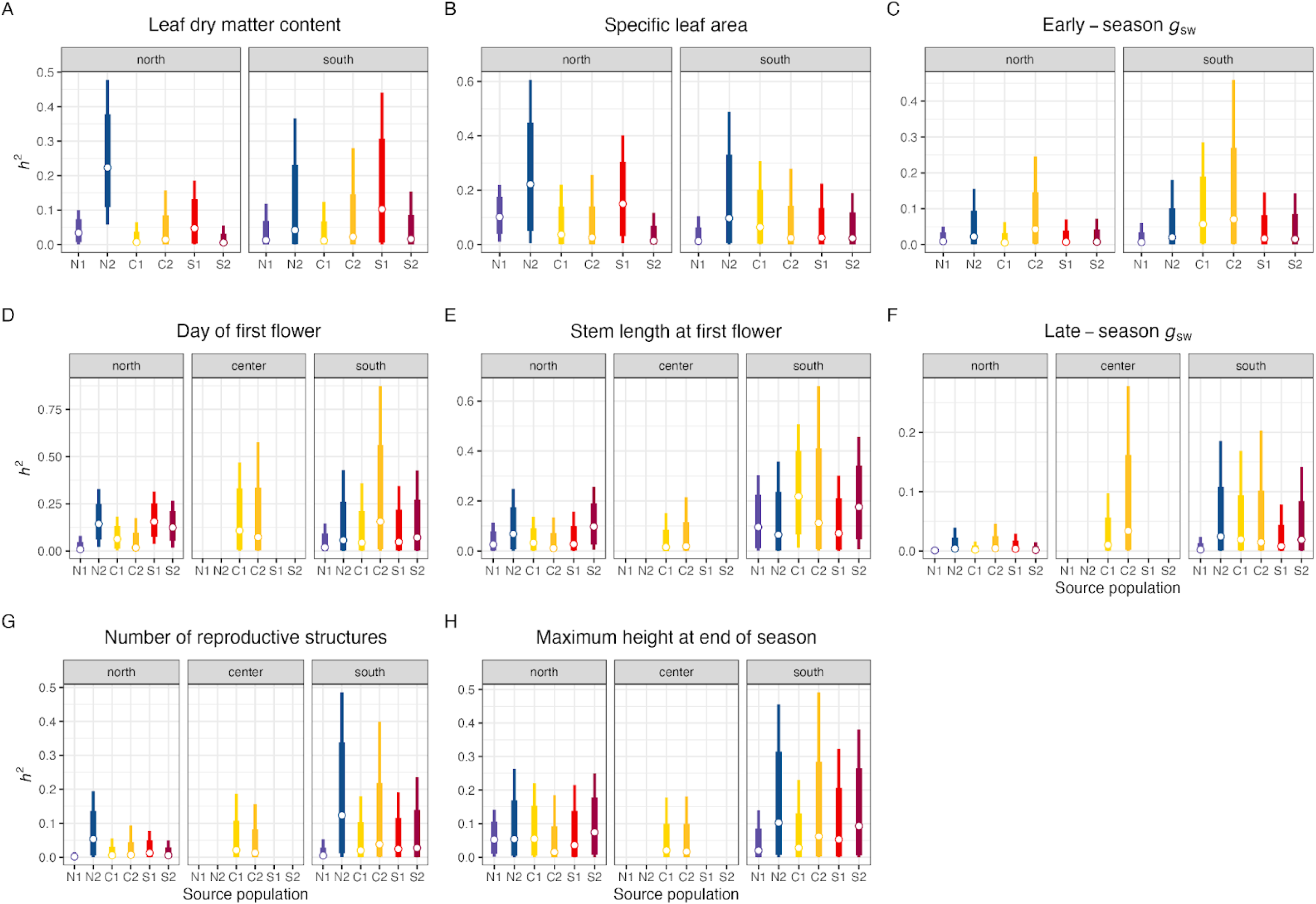
Narrow-sense heritability (*h^2^*) in traits for the ancestral cohort of each population in each garden. White circles represent median values, thick lines are 80% confidence intervals, and thin lines are 95% confidence intervals. Due to limited space in the central gardens, we were only able to estimate *h^2^* for the central populations, and logistical constraints prevented us from collecting data on leaf traits and early-season stomatal conductance (*g_sw_*) in the central garden.

### Do evolutionary responses in traits differ among populations or gardens (H1, H2)?

Overall, the difference between ancestral and descendant cohorts was small relative to the total trait variation in all populations and gardens (Table S10, Fig. 6). This result suggests that evolutionary response to drought contributes little to trait expression in semi-natural conditions. In the northern garden, descendants of N1 and S1 populations flowered later than their corresponding ancestors (Table S10, Fig. 6). In the central garden, the C2 population evolved an increase in late-season *g*_sw_, whereas the S1 population evolved reduced late-season *g*_sw_ (Table S10, Fig. 6). In the southern garden, the S1 population evolved an increase in *LDMC*, N1 evolved lower early-season *g*_sw_, S2 evolved a smaller stem height at first flower, N2 and S1 evolved reduced late-season *g*_sw_, and C2 evolved an increase in late-season *g*_sw_ from ancestors to descendants (Table S10, Fig. 6). All other populations in each garden did not show evolutionary shifts in any other trait (95% CIs overlapped 0; Table S10, Fig. 6).

**Figure 6.**
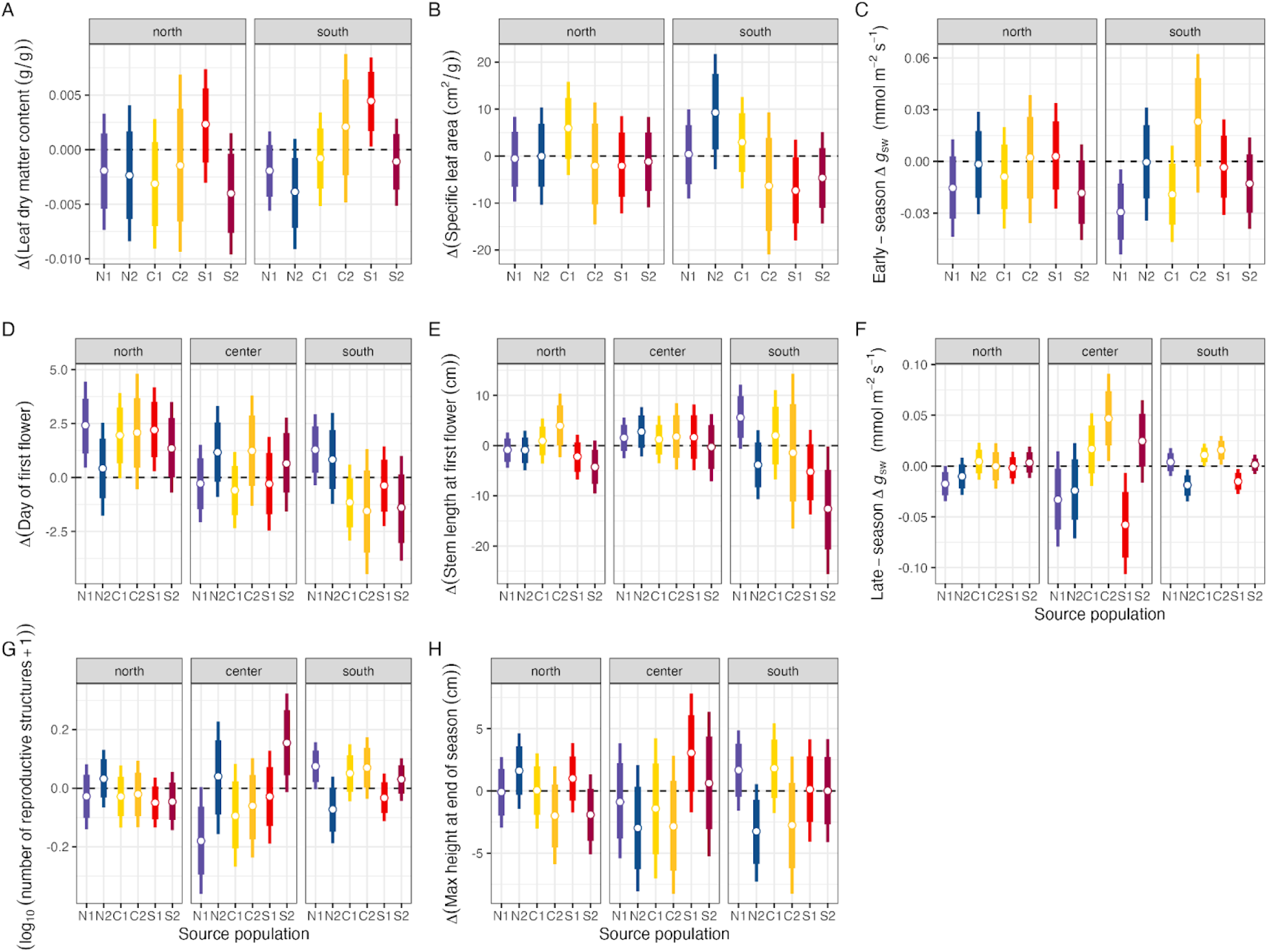
Model-predicted difference in each trait between descendants and ancestors of each population in each garden. White circles represent median values, thick lines are 80% confidence intervals, and thin lines are 95% confidence intervals. Dashed line corresponds to no change between descendants and ancestors. Due to logistical constraints, we did not collect data on leaf traits and early-season stomatal conductance (*g_sw_*) in the central garden.

## Discussion

The probability that evolution will rescue declining populations following extreme climate events can vary across species’ ranges, yet little is known about rangewide variation in adaptive potential. Despite the growing use of resurrection studies to evaluate evolutionary responses to recent climate change (Bishop *et al*. 2023; Dickman *et al*. 2019; Franks *et al*. 2007; Hamann *et al*. 2018; Kooyers *et al*. 2021; Kuester *et al*. 2016; Rauschkolb *et al*. 2023), to our knowledge this work represents the first to examine rates of trait evolution for central and edge populations in multiple field environments across a species’ range (Hendry *et al*. 2018; Shaw & Etterson 2012). Although we detected evolutionary differences in a small number of traits between ancestors and descendants of some populations in some gardens, these responses were small in magnitude and did not follow expectations from range limit theory (H1). Nonetheless, the few observed evolutionary shifts resulted in environmentally-dependent changes in trait expression under field conditions, whereby the magnitude of trait change between ancestors and descendants varied among experimental gardens (H2). Due to the relatively low and imprecise estimates of heritability and the limited evolutionary shifts in traits, there was no support for the prediction from the breeder’s equation that populations with greater trait heritabilities would show greater absolute magnitudes of evolutionary change in traits. Similarly, although populations experienced different magnitudes of drought stress, precipitation anomalies did not explain trait differences between ancestors and descendants, also because of limited evolutionary responses. Together, these preliminary results from a single growing season suggest that the expression of genetic variation, and thus traits, depend on the environmental context, and that environmental variability in field conditions may mask the genetic variation that is often expressed in greenhouse settings (Conner *et al*. 2003). Below, we discuss the potential mechanisms underlying our main findings and the implications for our understanding of evolutionary rescue across species’ ranges.

### Backdrop of local maladaptation

We conducted this study against the backdrop of local maladaptation, whereby nonlocal genotypes outperformed local genotypes (Brady *et al*. 2019), at the leading range edge (Bontrager *et al*. 2021) and range center. Specifically, southern populations had greater fitness than central populations in the central garden and both southern and central populations had greater fitness than northern populations in the northern garden (Fig. 3g). This context of maladapted northern and central populations, potentially due to climate change, is important for studying which traits underlie these fitness advantages in populations originating from warmer climates–e.g., whether these populations have evolved drought escape or avoidance strategies. Several common garden and reciprocal transplant studies have shown local maladaptation or lags in adaptation to current climate (Albano *et al*. 2024; Anderson & Wadgymar 2020; Bradley St Clair & Howe 2007; Gorton *et al*. 2022; Wilczek *et al*. 2014), including a study of five *M. cardinalis* populations transplanted into a northern and southern common garden (Muir and Angert, *unpublished data*). Surprisingly, when we transplanted northern and central populations southward into an environment with greater climatic moisture deficit (Table S1), descendants of a northern population tended to have reduced fitness in the central garden, and descendants of the same northern population and both central populations showed a trend of elevated fitness relative to their ancestors in the southern garden (Fig. 6g). Though these trends are small in magnitude, this suggests that some populations could have the ability to evolve increased performance under a changing climate, especially on longer timescales.

### Plasticity and genetic differentiation in escape and avoidance traits

The direction of plastic, but not genetic, changes in traits is consistent with the hypothesis that *Mimulus cardinalis* escapes rather than avoids drought. In the most xeric, southern garden, plants developed cheaper leaves (lower *LDMC* and higher *SLA*) and had higher early-season *g*_sw_ than counterparts in the northern garden. Plants in the central garden flowered slightly later and had higher late-season *g*_sw_ than at either other garden. The phenotypes at the central garden are difficult to interpret because the delays in transplanting precluded early-season measurements.

Furthermore, these plants were developmentally younger than plants at other gardens measured at the same part of the season, which may explain their later flowering time and higher late-season *g*_sw_. Within gardens, source populations from different regions exhibited similar average *SLA* and stomatal conductance, but showed stronger genetic differentiation in *LDMC*, first flower date, and size at first flower (Fig. 3). Lack of genetic differentiation among populations in some traits in the experimental gardens, likely resulting from greater environmental variation, contrasts with multiple observations of genetic differences in similar traits measured in controlled environments (Angert *et al*. 2011; Branch *et al*. 2024; Muir *et al*. 2022; Muir & Angert 2017; Nelson *et al*. 2021; Sheth & Angert 2016). However, genotype-by-environment interactions are likely important too because the phenotypic and fitness effects of alleles in the field can be difficult to predict from studies in controlled environments (El-Soda *et al*. 2014). The sample size of our experiment was sufficient to detect differences among populations comparable to that observed in controlled experiments, despite greater environmental variance.

### Limited trait evolution between ancestors and descendants

Though several resurrection studies, including some with *M. cardinalis*, have documented rapid evolutionary responses in traits associated with drought adaptation (Anstett *et al*. 2021; Branch *et al*. 2024; Christie *et al*. 2023; Dickman *et al*. 2019; Franks 2011; Franks *et al*. 2007; Franks & Weis 2008; Hamann *et al*. 2018; Kooyers *et al*. 2021; Rauschkolb *et al*. 2022), we detected limited evolutionary responses in any of our focal escape and avoidance traits. This overall lack of response precluded our ability to test hypotheses about the roles of genetic variation and the strength of selection in determining the magnitude of evolutionary response. Given that most point estimates of heritability for ancestral cohorts were close to 0 (*h^2^* < 0.1 for most population × trait × garden combinations), populations could have lacked sufficient genetic variation in escape and avoidance traits. Similar to several other studies (Conner *et al*. 2003; Geber & Griffen 2003; Hoffmann *et al*. 2017), the low heritabilities we estimated in the field contrast with high heritabilities of some traits for some *M. cardinalis* populations in more controlled greenhouse settings (Muir *et al*. 2022; Nelson *et al*. 2021; Sheth & Angert 2016). For instance, an artificial selection study involving the ancestral cohort of the same populations of *M. cardinalis* found that the two southern-edge populations rapidly responded to selection for early and late flowering (Sheth & Angert 2016). Moreover, the imprecision around heritability estimates, possibly due to environmental variation (e.g., patches of weeds, uneven irrigation due to topography) within each garden prevented us from detecting among-population variation in heritability, a requirement for robustly testing the hypothesis that the magnitude of trait evolution is proportional to standing genetic variation for that trait. Notably, heritability estimates were most precise in the northern garden (Fig. 5), potentially due to lower environmental variation than the southern and central gardens. Bemmels and Anderson (2019) similarly documented low and imprecise estimates of genetic variation for most traits examined in a field study of the perennial forb, *Boechera stricta*. Despite the challenges associated with estimating quantitative genetic parameters precisely under field conditions, combining these field estimates with measures of phenotypic selection is essential for making and testing predictions about evolutionary rescue in the wild (Campbell 1996; Grant & Grant 1995).

Limited evolution in some traits could also be explained by a lack of strong, directional selection on these traits. For instance, in the southern garden, which represents the conditions with the greatest climatic moisture deficit in the study (Table S1), *SLA* and *LDMC* were under weak, stabilizing and directional selection, respectively, and there was evidence for stabilizing selection in size at first flower and at the end of the growing season (Fig. 4). Since the strength and mode of selection can vary among years (Siepielski *et al*. 2009) and we have only estimated selection from a single growing season, these estimates cannot fully capture patterns of selection that occurred during the extreme drought event. Because there were only three instances in which we detected nonzero evolutionary responses of populations transplanted southward into environments with higher climatic moisture deficit, we had limited ability to evaluate whether temporal evolutionary responses to recent drought mirror spatial patterns of selection.

Inconsistent with an adaptive response to warmer, drier environments, in two of the three cases, the direction of evolutionary response countered the direction of selection. Specifically, in the southern garden, N1 evolved lower early-season *g*_sw_, N2 evolved lower late-season *g*_sw_, and C2 evolved higher late-season *g*_sw_. For the two northern populations, evolutionary responses opposed the direction of selection for increased early- and late-season *g*_sw_ in the southern garden. Conversely, for the C2 population, the evolutionary response was consistent with selection for increased late-season *g*_sw_ in the southern garden (Figs. 4, 6). Additional examinations of evolutionary responses reveal similar patterns–for example, descendants of one northern population flowered later than ancestors even though selection in all gardens, and especially in the southern garden, favored early flowering (Figs. 4, 6). Coincidentally, we established the experimental gardens for this study during a year in which California experienced historic precipitation that ended a multi-year drought (DeFlorio *et al*. 2024). These anomalous conditions could have led to substantially different patterns of selection than what these populations experienced during the extreme drought event, and played a role in decoupling spatial and temporal responses to climate. Even so, populations varied considerably in the magnitude of drought they experienced (Fig. 2), and theory and microcosm experiments suggest that evolutionary rescue is most likely when environmental change is gradual (Carlson *et al*. 2014; Gonzalez & Bell 2013; Lindsey *et al*. 2013). Yet, the influence of the rate of environmental change on the likelihood of evolutionary rescue in natural populations of longer-lived, sexually reproducing organisms is poorly understood (Bell 2017; Carlson *et al*. 2014). All of the greenhouse and growth chamber studies involving the same populations of *M. cardinalis* as our field study similarly found an overall lack of evolutionary response in traits associated with thermal adaptation (Wooliver et al. 2020) and phenology (Vtipil & Sheth 2020). In contrast, another study examining the evolution of drought avoidance and escape traits in *M. cardinalis* with greater spatial and temporal sampling found that southern populations showed rapid evolutionary shifts from avoidance to escape in just 7 years (Anstett *et al*. 2021), discounting the possibility that populations of this perennial species have simply not had enough time to adapt to recent changes in climate.

### Environmental dependence of the expression of genetic variation

Consistent with studies showing that the expression of genetic differences within and among populations and cohorts can vary across environments (Bemmels & Anderson 2019; Karitter *et al*. 2023), we found that differences in trait expression between ancestors and descendants often varied among gardens (Fig. 6), as did the precision of heritability estimates (Fig. 5). A comparison of trait differences between ancestors and descendants of the perennial herb *Leontodon hispidus* measured in greenhouse, growth chamber, and outdoor settings revealed that the magnitude of trait difference varied across environments, and there were even some differences that were only detectable in a subset of environments (Karitter *et al*. 2023).

Similarly, resurrection studies of the annual mustard, *Brassica rapa*, showed that the magnitude of evolutionary change between ancestors and descendants varied with watering treatment, but the direction of change was generally consistent (Hamann *et al*. 2018; Johnson *et al*. 2022). In contrast, we found that the direction of trait change sometimes varied among gardens. For example, central and southern populations showed a trend of evolving delayed flowering from ancestors to descendants in the northern garden, but advanced flowering in the southern garden (Fig. 6d), which could have downstream ecological consequences such as decoupling species interactions (Olliff-Yang & Mesler 2018; Stemkovski *et al*. 2020). Though populations may adapt to drought via the evolution of escape or avoidance traits, drought stress could alternatively mask the expression of trait evolution (Bemmels & Anderson 2019), but further work is needed to understand why trait evolution between ancestors and descendants differed among gardens. Similarly, trait heritabilities of populations transplanted into warmer, drier climates could either increase or decrease in these novel environments (Bemmels & Anderson 2019; Hoffmann & Merilä 1999). Our point estimates of heritability did not clearly differ among gardens, but estimates were generally more precise in the northern than the southern garden (Fig. 5). Thus, we do not have clear evidence pointing towards populations having lower or higher heritabilities when transplanted into non-local gardens.

### Caveats

There are several caveats that hinder the inferences we can draw about evolutionary rescue from this work. First, since we have presented results from one growing season of a perennial plant, the data from this study represent short-term, early-stage responses and do not incorporate the fitness components of germination success and seedling survival. Given that our findings already highlight the environmental dependence of phenotypic selection, quantitative genetic variation, and the expression of traits and fitness, accounting for interannual variation in these parameters is crucial to fully understand evolutionary responses to the recent drought event. Future years will reveal how these early-stage patterns relate to longer-term fitness.

Ultimately, a complete test of evolutionary rescue requires long-term data on the dynamics of natural populations, and this is the focus of ongoing work (Anstett *et al*. 2024). Second, various aspects of the experimental gardens do not perfectly capture natural riparian environments in terms of shade, soil moisture, inter- and intraspecific competition, and other biotic interactions. There were also other confounding factors unrelated to climate, such as variation in transplant date among gardens (Table S1), that could have influenced the focal phenotypes and fitness proxies. Despite these limitations, the experimental garden approach narrows the gap between greenhouse studies and natural field environments by allowing us to isolate the effects of broad climatic differences on traits and fitness. Third, even if populations harbor sufficient genetic variation in a single trait, they may lack appropriate genetic variation in multiple traits important for responding to natural selection, and genetic trade-offs between traits could constrain multivariate responses to selection (Etterson & Shaw 2001; Gomulkiewicz & Houle 2009; Walsh & Blows 2009). Here, we focused exclusively on each trait independently, but future work with additional years of data will examine multivariate evolutionary responses and the **G** matrix in the context of the multivariate breeder’s equation (Chantepie *et al*. 2024; Conner *et al*. 2003; Henry & Stinchcombe 2023; Lande 1979; Lande & Arnold 1983). With additional years of trait and fitness data, we will also investigate the covariance between traits and fitness based on the Robertson-Price identity (Bemmels & Anderson 2019; Etterson & Shaw 2001; Morrissey *et al*. 2010), and the fundamental theorem of natural selection, which predicts the change in mean fitness from one generation to the next as the ratio of additive genetic variance for fitness to mean fitness (Bonnet *et al*. 2022; Kulbaba *et al*. 2019; Peschel & Shaw 2024; Shaw 2019; Shaw & Etterson 2012; Torres-Martínez *et al*. 2019). Fourth, previous work suggests that a small proportion of *M. cardinalis* seeds can remain viable in the seed bank beyond the first year (Angert 2006a), potentially slowing the pace of evolutionary change (Benning *et al*. 2023; Hairston Jr & De Stasio Jr 1988). Given that other resurrection studies with *M. cardinalis* conducted across similar years have found evidence for evolutionary change (Anstett *et al*. 2021; Branch *et al*. 2024), and that a demographic study of *M. cardinalis* that overlapped with the period of historic drought found that seeds did not remain viable in the seed bank beyond the first year (Sheth & Angert 2018), we consider this possibility to be unlikely. Finally, if rapid evolution is insufficient to rescue populations, adaptive plasticity may provide a critical mechanism for persistence, yet few studies explore how adaptive plasticity and local adaptation may work together to rescue populations from severe drought (Chevin *et al*. 2013; Scheiner *et al*. 2017; Snell-Rood *et al*. 2018). Including simulations of warm, dry conditions within each garden (e.g., via drought treatments and/or open-top warming chambers) would permit robust studies of the role of phenotypic plasticity in rescuing declining populations (Peschel *et al*. 2021). Ongoing work is investigating whether populations have evolved changes in plasticity in escape or avoidance traits after the historic drought, a question that remains unresolved in resurrection studies, particularly in field settings (Anstett *et al*. 2021; Franks 2011; Hamann *et al*. 2018; Johnson *et al*. 2022).

### Conclusions

To our knowledge, this work represents the first resurrection study to quantify selection, trait heritabilities, and rates of trait evolution for populations across a species’ range in field settings, including near range limits. We build upon the current understanding of evolutionary rescue, which is based primarily on simulation and laboratory studies, by assessing the potential for evolutionary rescue across the geographic range of *Mimulus cardinalis* against the backdrop of local maladaptation in central and leading-edge populations. We also investigate the suite of traits that determine the likelihood of adaptation or maladaptation to recent climate change in a perennial plant, which are often understudied despite their prevalence in nature, due to longer generation times leading to potentially slower evolutionary responses. Though we detected limited evolutionary change between ancestors and descendants over a 7-year period, future seed collections from the same populations will enable investigations of long-term evolutionary responses to climate change in a perennial plant species. Nonetheless, understanding the causes of evolutionary stasis in natural populations is a longstanding challenge in evolutionary ecology (Merilä *et al*. 2001). Investigating whether range-edge populations have evolved in response to recent drought is crucial for predicting range shifts under climate change, yet most range shift forecasts ignore the potential for evolution and likely overestimate vulnerability to climate change (Anderson *et al*. 2025; Bush *et al*. 2016). Ultimately, parameter estimates from these and follow-up experiments will allow ecologists to account for real-time evolutionary responses in their forecasts of range shifts.

## Acknowledgements

We thank Nathanael Larson, Lauryn Cabral, Genevieve Triplett, Gabe Bradshaw, Brooke Crose, Rachel Muir, Sarah Erskine, Jeremy Collings, Bryn Callie, Austyn Tavernier, Anna Easton, and several short-term field technicians who helped set up the experiment and collect data. We are grateful to greenhouse staff at the University Oregon, San Diego State University, and the University of California, Merced for support with plant growth facilities. James Bourdon, Alexander Powell, Jacob Keeton, and Pablo Bryant at the Santa Margarita Ecological Reserve, Mieko Aoki and Hal Cushman at the Friends of Buford Park & Mt. Pisgah Native Plant Nursery, and Joanna Clines from the Sierra National Forest provided permission and tremendous logistical and infrastructural support for this project. Jeff Conner, Ruth Shaw, and Amy Angert contributed advice related to experimental design and data collection. We thank Cuyamaca Rancho State Park, the Bureau of Land Management, and Yosemite National Park for granting collecting permits. Crosses that produced the refresher generation were performed by A. Coughlin, R. Wooliver, E. Vtipilthorpe, and A. Wiegmann, and C. Payst, S. Correa, C. Laufenberg, and E. Wilson assisted with data entry and seed preparation. This work was supported by National Science Foundation projects DEB-2131815 (SNS), 2131816 (JD), 2131817 (CDM), 2131818 (JPS), and 2131819 (LF-R), a Research Capacity Fund (HATCH) project award no. 7002993 from the U.S. Department of Agriculture’s National Institute of Food and Agriculture (SNS), and a North Carolina State University Goodnight Early Career Innovators Award (SNS). This work was performed (in part) at the University of California Natural Reserve System Boyd Deep Canyon Desert Research Center (Reserve DOI: <u>10.21973/N3V66D</u>) and Sierra Nevada Research Station – Yosemite Field Station (Reserve DOI: <u>10.21973/N3V36C</u>). Associate Editor Lauren Buckley, John Benning, and two anonymous reviewers provided invaluable feedback on earlier drafts of this work. SNS is grateful to the Parmar, Patel, Randle/Paul, Saraiya/Nagar and Bishop families for their support along the way.

## Author contributions

SNS, CDM, JD, LF-R, and JPS conceptualized and designed the study, and obtained funding to support the work. KK, MM, EJC, and DW helped develop protocols for data collection. All authors except LJA collected the data. SNS, CDM, and LJA analyzed the data and wrote the original draft of the manuscript. SNS, LJA, CDM, and LF-R contributed to conceptual and data visualization. LF-R, JPS, JW, EJC, and JD edited the manuscript. All authors reviewed and approved the manuscript.

## Dryad link with data and code

http://datadryad.org/stash/share/QuYUWKTibcxWQqbJ3_G4Q-28Kma51suAEoxH18ROj58

