## Supplementary Materials for "Evolutionary responses to historic drought across the range of scarlet monkeyflower"

**Supplementary Material**

**Supplementary Methods and Results**

*Validation of fitness proxies through subset measurements*

In the main analysis of this study, we used proxies for plant size and sexual reproduction in order to avoid potential challenges in data collection. Namely, we used maximum plant height, measured in centimeters as the length of the tallest stem, to represent total vegetative growth of each plant, and counts of reproductive structures (buds, flowers, fruits, and pedicels) on three stems per plant (or fewer if a plant had only one or two reproductive stems) to estimate the total number of reproductive structures produced by each plant. For plants with three or more reproductive stems, we estimated the total number of reproductive structures for the plant as: *# of structures on the stem with the maximum # of structures + (sum of the # of structures on the 2 other stems)/2) * (# of reproductive branches − 1)*. For plants with two reproductive stems, the total number of reproductive structures was estimated as: *# of structures on the stem with the maximum # of structures + (number of structures on the other stem * (# of reproductive branches − 1))*. For plants with one reproductive stem, the total number of reproductive structures is simply the number of structures on that stem. We also recorded each plant's survival and probability of flowering, and assigned plants that never flowered a value of zero for the number of reproductive structures, whether they survived the growing season or not.

We validated our use of proxies to estimate plant size and fitness by conducting a more extensive data collection on a subset of 100 plants per garden and performing correlations between our chosen proxies and other variables that may more accurately reflect plant growth and reproduction. The subset of plants was selected haphazardly within each garden, aiming for approximately equal representation among populations and cohorts. For these plants, we measured the total number of fruits that successfully set seed and total aboveground dry biomass (g), along with the total length of all stems (cm), which was estimated through the following calculations. For plants with greater than 3 stems, total stem length was estimated as: *length of the longest stem + ((length of a 2nd stem + length of a 3rd stem)/2) * (number of branches − 1)*, with the 2nd and 3rd stems being chosen in a manner that was representative of all available stems on a given plant. If a plant had 3 or fewer stems, the lengths of each measured stem were summed to calculate total stem length. We then validated our estimates of total number of reproductive structures per plant in the main analysis by performing correlations between the estimated total number of reproductive structures (regardless of successful seed production; Total # of RS) and the total number of successful fruits (those that produced viable seed; Total # of SF), Total # of SF and dry biomass, and Total # of RS and dry biomass for the subset of plants within each garden. We also validated our use of maximum plant height in the main analysis by performing correlations between maximum height and total stem length, maximum height and biomass, and total stem length and biomass for the subset of plants within each garden. Pearson's correlation coefficients ($r$) and their associated $P$ values were generated using the *cor* and *cor.test* functions, respectively, in R version 4.5.0 (R Core Team, 2025). In all three experimental gardens, each of these correlations was positive and statistically significant ($P < 0.05$; Table S3; Figs. S1, S2), providing validation for our use of proxy measurements in the main analysis of this study.

*Estimates of $h^2$ from brms and MCMCglmm sub-models*

As a robustness check, we compare the global estimates of the medians and 95% confidence intervals for $h^2$ from brms to mini-models from brms v. 2.22.0 and MCMCglmm v. 2.36 (Hadfield 2010). Specifically, we focus on a single trait, flowering time, because we have data for this trait in all gardens, and we report these estimates using the ancestral cohorts of "local" populations within each garden (e.g., populations N1 and N2 in the northern garden, populations C1 and C2 in the central garden, and populations S1 and S2 in the southern garden). Thus, these sub-models include flowering time as the response variable, and the random effects of sire, dam nested within sire, and garden block, with a separate model for each "local" population in each garden. The fixed and random effect model structure was identical between sub-models, but MCMCglmm uses different default weakly informative prior distributions. The MCMCglmm model ran for 10,000 iterations with about 3,000 burnin iterations on a single chain. As with the brms model, most parameters converged ($\widehat{R} < 1.05$) and were adequately sampled (ESS > 500). However, some constrained parameters and random effect coefficients did not quite meet these convergence criteria (see further explanation in main text). Overall, estimates of $h^2$ were similar based on visual inspection, though the lower 95% CI and median $h^2$ estimates for the S1 and S2 populations in the southern garden were slightly higher for both subset models relative to the global model (Table S5, Figure S3).

*Visualization of G x E*

To visualize the nature of genotype-by-environment interactions (G X E), we plot breeding values for each population across pairs of gardens (as in Sheth et al. 2018), using flowering time as an example. Overall, breeding values were positively correlated within each population in each garden, but the strength of these correlations varies among populations and gardens (Fig. S4).

**Supplementary Tables [note: Tables S5 - S9 are provided as .csv files zipped together]**

**Table S1**. Descriptive information for each of the common garden locations used in the field experiment. Climate data are derived from the web version of ClimateNA v. 4.60 (Wang et al. 2016).

| | North | Center | South |
|---|---|---|---|
| Latitude | 44°01’05” | 37°22’46” N | 33°26’18” N |
| Longitude | 122°59’15” | 119°37’44” W | 117°10’35” W |
| Elevation (m) | 138 | 967 | 262 |
| Total Number of Plants | 5468 | 3354 | 5468 |
| Total Number of Blocks | 10 | 11 | 10 |
| Garden Area | 744 $m^2$ | 499 $m^2$ | 791 $m^2$ |
| Planting Dates (Greenhouse) | 02/27/23 - 03/03/23 | 04/24/23 - 05/01/23 | 03/21/23 - 03/27/23 |
| Transplanting Dates (Field) | 04/06/23 - 04/26/23 | 06/08/23 - 06/17/23 | 05/10/23 - 05/26/23 |
| Mean annual temperature | 12.0 °C | 9.9 °C | 17.3 °C |
| Winter precipitation | 336 mm | 1337 mm | 490 mm |
| Climatic moisture deficit | 574 mm | 920 mm | 878 mm |

**Table S2**. The number of unique sires and dams from each focal population transplanted into each of the three common garden locations.

| Garden | N1 | | N2 | | C1 | | C2 | | S1 | | S2 | |
|---|---|---|---|---|---|---|---|---|---|---|---|---|
| | sires | dams | sires | dams | sires | dams | sires | dams | sires | dams | sires | dams |
| north | 8 | 36 | 16 | 72 | 28 | 132 | 9 | 44 | 33 | 157 | 32 | 158 |
| center | 8 | 36 | 16 | 68 | 28 | 133 | 9 | 44 | 25 | 99 | 25 | 100 |
| south | 8 | 36 | 16 | 70 | 28 | 133 | 9 | 44 | 33 | 161 | 32 | 157 |

**Table S3**. Pearson's correlations (*r*) between proxies of fitness of *M. cardinalis* plants in each of the three common gardens (north, center, and south). Proxies used in the main analysis of the paper included maximum plant height and the total number of reproductive structures per plant (Total # of RS). Variables used for validation of these proxies include the total number of successful fruits (Total # of SF), plant dry biomass, and total stem length of the entire plant. Significant correlations ($p < 0.05$) are denoted in bold.

| | x = Total # of RS<br>y = Total # of SF | | x = Biomass<br>y = Total # of SF | | x = Biomass<br>y = Total # of RS | | x = Max Height<br>y = Total Stem Length | | x = Max Height<br>y = Biomass | | x = Total Stem Length<br>y = Biomass | |
|---|---|---|---|---|---|---|---|---|---|---|---|---|
| Garden | *r* | *p* | *r* | *p* | *r* | *p* | *r* | *p* | *r* | *p* | *r* | *p* |
| north | **0.981** | **< 0.001** | **0.826** | **< 0.001** | **0.842** | **< 0.001** | **0.551** | **< 0.001** | **0.611** | **< 0.001** | **0.763** | **0.014** |
| center | **0.828** | **< 0.001** | **0.859** | **0.001** | **0.895** | **< 0.001** | **0.448** | **< 0.001** | **0.630** | **< 0.001** | **0.390** | **< 0.001** |
| south | **0.987** | **< 0.001** | **0.861** | **< 0.001** | **0.854** | **< 0.001** | **0.327** | **0.012** | **0.698** | **< 0.001** | **0.429** | **< 0.001** |

**Table S4**. Sample sizes for each of the eight response variables involved in the study: leaf dry matter content (*LDMC*), specific leaf area (*SLA*), early-season stomatal conductance (early $g_{sw}$), day of first flower, total stem length at time of first flower, late-season stomatal conductance (late $g_{sw}$), total number of reproductive structures (Total # of RS), and maximum plant height. Sample sizes are divided by garden, along with source population from the northern range edge (N1, N2), latitudinal range center (C1, C2), and southern range edge (S1, or S2) and cohort (2010 or 2017). Full gardens, blocks ("B"), or rows ("R") within gardens that were excluded from data collection for particular response variables are also identified.

| north | | | | | | | | | | | | | |
|---|---|---|---|---|---|---|---|---|---|---|---|---|---|
| | | N1 | | N2 | | C1 | | C2 | | S1 | | S2 | |
| Trait | Blocks/Rows Excluded | 2010 | 2017 | 2010 | 2017 | 2010 | 2017 | 2010 | 2017 | 2010 | 2017 | 2010 | 2017 |
| *LDMC* | None | 262 | 149 | 492 | 135 | 609 | 147 | 342 | 123 | 639 | 150 | 663 | 190 |
| *SLA* | None | 261 | 150 | 491 | 135 | 607 | 147 | 340 | 122 | 642 | 149 | 657 | 189 |
| Early $g_{sw}$ | None | 268 | 150 | 501 | 138 | 616 | 148 | 342 | 123 | 649 | 153 | 671 | 195 |
| Day of First Flower | None | 333 | 191 | 664 | 188 | 779 | 222 | 411 | 166 | 880 | 216 | 811 | 271 |
| Total Stem Length | None | 333 | 191 | 664 | 188 | 779 | 222 | 411 | 166 | 879 | 216 | 811 | 271 |
| Late $g_{sw}$ | None | 298 | 179 | 597 | 173 | 714 | 205 | 382 | 160 | 850 | 208 | 755 | 253 |
| Total # of RS | B5, B6, R701-702 | 243 | 155 | 480 | 142 | 560 | 171 | 306 | 123 | 661 | 157 | 633 | 215 |
| Max Height | B5, B6, R701-702 | 237 | 151 | 481 | 144 | 565 | 166 | 305 | 123 | 668 | 160 | 622 | 209 |

| center | | | | | | | | | | | | | |
|---|---|---|---|---|---|---|---|---|---|---|---|---|---|
| | | N1 | | N2 | | C1 | | C2 | | S1 | | S2 | |
| Trait | Blocks/Rows Excluded | 2010 | 2017 | 2010 | 2017 | 2010 | 2017 | 2010 | 2017 | 2010 | 2017 | 2010 | 2017 |
| *LDMC* | All | 0 | 0 | 0 | 0 | 0 | 0 | 0 | 0 | 0 | 0 | 0 | 0 |
| *SLA* | All | 0 | 0 | 0 | 0 | 0 | 0 | 0 | 0 | 0 | 0 | 0 | 0 |
| Early $g_{sw}$ | All | 0 | 0 | 0 | 0 | 0 | 0 | 0 | 0 | 0 | 0 | 0 | 0 |
| Day of First Flower | None | 167 | 173 | 199 | 174 | 719 | 185 | 398 | 243 | 184 | 182 | 178 | 244 |
| Total Stem Length | None | 172 | 178 | 203 | 180 | 739 | 188 | 412 | 250 | 190 | 188 | 186 | 257 |
| Late $g_{sw}$ | B8 | 126 | 142 | 148 | 150 | 581 | 142 | 324 | 198 | 163 | 160 | 165 | 226 |
| Total # of RS | B8, B11 | 129 | 139 | 145 | 146 | 538 | 130 | 287 | 182 | 158 | 145 | 158 | 199 |

| | | | | | | | | | | | | | |
|---|---|---|---|---|---|---|---|---|---|---|---|---|---|
| Max Height | B8, B11 | 125 | 130 | 140 | 144 | 519 | 121 | 288 | 179 | 154 | 144 | 149 | 197 |

| south | | | | | | | | | | | | | |
|---|---|---|---|---|---|---|---|---|---|---|---|---|---|
| | | N1 | | N2 | | C1 | | C2 | | S1 | | S2 | |
| Trait | Blocks/Rows Excluded | 2010 | 2017 | 2010 | 2017 | 2010 | 2017 | 2010 | 2017 | 2010 | 2017 | 2010 | 2017 |
| *LDMC* | None | 301 | 202 | 573 | 194 | 779 | 223 | 405 | 179 | 863 | 207 | 904 | 283 |
| *SLA* | None | 299 | 202 | 562 | 192 | 766 | 219 | 401 | 175 | 848 | 200 | 884 | 282 |
| Early $g_{sw}$ | None | 305 | 208 | 575 | 193 | 785 | 226 | 410 | 180 | 867 | 210 | 910 | 289 |
| Day of First Flower | None | 332 | 225 | 648 | 215 | 809 | 230 | 421 | 183 | 901 | 221 | 928 | 293 |
| Total Stem Length | None | 332 | 225 | 648 | 214 | 809 | 229 | 421 | 182 | 898 | 221 | 926 | 293 |
| Late $g_{sw}$ | B3, B9 | 205 | 144 | 445 | 147 | 605 | 162 | 309 | 137 | 687 | 172 | 713 | 234 |
| Total # of RS | B3, B8, B9, R501-502 | 195 | 132 | 389 | 127 | 487 | 130 | 251 | 108 | 573 | 141 | 563 | 179 |
| Max Height | B3, B8, B9, R501-502 | 197 | 134 | 391 | 125 | 482 | 131 | 253 | 110 | 572 | 140 | 566 | 182 |

**Table S5 (.csv file)**. Estimates of medians and 95% CIs (“lower” and “upper” columns) for narrow-sense heritability ($h^2$) for flowering time for each population in its local garden (i.e., populations N1 and N2 in northern garden, C1 and C2 in central garden, and S1 and S2 in southern garden). Estimates are derived from three methods: 1) a global model including data from all cohorts, populations, and gardens using the brms package (as in Table S8 and Fig. 5), and models built with data from only the ancestral cohort, separately for each population in each garden using 2) brms, and 3) MCMCglmm packages.

**Table S6 (.csv file)**. Medians and 95% CIs (“lower” and “upper” columns) for each trait of each cohort of each population in each garden. Traits include leaf dry matter content (“ldmc”), specific leaf area (“sla”), early-season stomatal conductance (“gsw_early”), day of first flower (“first_flower_doy”, total stem length at first flower (“totalStemLen”), late-season stomatal conductance (“gsw_late”), $\log_{10}$-transformed total number of reproductive structures (“log_totalRS”), and maximum plant height (“maxHeight”). Traits are summarized by garden, along with source population from the northern range edge (N1, N2), latitudinal range center (C1, C2), and southern range edge (S1, or S2) and cohort (2010 or 2017).

**Table S7 (.csv file)**. Medians and 95% CIs (“lower” and “upper” columns) for each selection coefficient of each trait in each garden. Regression coefficients include the intercept $b_0$, the linear coefficient, $b_1$, and the quadratic coefficient, $b_2$. The quadratic coefficients from regressions were doubled to obtain quadratic selection gradients (“quad_sel_grad” column; Stinchcombe et al. 2008). Traits include leaf dry matter content (“ldmc”), specific leaf area (“sla”), early-season stomatal conductance (“gsw_early”), day of first flower ("first_flower_doy", total stem length at first flower ("totalStemLen"), late-season stomatal conductance ("gsw_late"), and maximum plant height ("maxHeight").

**Table S8 (.csv file)**. Medians and 95% CIs ("lower" and "upper" columns) for narrow-sense heritability ($h^2$), additive genetic variance ($V_A$), and environmental variance ($V_E$) of each trait for the ancestral (2010) cohort of each population in each garden. Traits include leaf dry matter content ("ldmc"), specific leaf area ("sla"), early-season stomatal conductance ("gsw_early"), day of first flower ("first_flower_doy", total stem length at first flower ("totalStemLen"), late-season stomatal conductance ("gsw_late"), $log_{10}$-transformed total number of reproductive structures ("log_totalRS"), and maximum plant height ("maxHeight"). Quantitative genetic parameters are summarized by garden, along with source population from the northern range edge (N1, N2), latitudinal range center (C1, C2), and southern range edge (S1, or S2).

**Table S9 (.csv file)**. Model selection table comparing a suite of models of additive genetic variance ($V_A$ ) for each trait. Because sample sizes only permitted robust estimation for the ancestral cohort of each population, all models also included a cohort effect to obtain separate estimates of $V_A$ for each cohort, but we only report $V_A$ estimates for ancestral cohorts. Columns include trait ("ldmc": leaf dry matter content, "sla": specific leaf area, "gsw-early": early-season stomatal conductance, "flowering_time": day of first flower, "totalStemLen": total stem length at time of first flower, "gsw-late": late-season stomatal conductance, "fitness": total number of reproductive structures, and "maxHeight": maximum plant height at the end of the growing season), model ("full": full model with separate $V_A$ for each population in each garden; "garden": model with separate $V_A$ for each garden but no differences in $V_A$ among populations; “population”: model with separate $V_A$ for each population but no differences in $V_A$ among gardens; and “minimal”: model with sire and dam effects removed, representing no $V_A$), looic: Bayesian leave-one-out criterion (*LOOIC*) scores, se_looic: standard errors for *LOOIC*, deltalooic: pairwise differences between models in *LOOIC* scores ($\Delta LOOIC$), and se_deltalooic: standard errors for $\Delta LOOIC$.

**Table S10 (.csv file)**. Median and 95% CIs (“lower” and “upper” columns) for evolutionary response (descendant trait value - ancestor trait value) in each trait for each population in each garden. Traits include leaf dry matter content (“ldmc”), specific leaf area (“sla”), early-season stomatal conductance (“gsw_early”), day of first flower (“first_flower_doy”, total stem length at first flower (“totalStemLen”), late-season stomatal conductance (“gsw_late”), $\log_{10}$-transformed total number of reproductive structures (“log_totalRS”), and maximum plant height (“maxHeight”). Evolutionary responses are summarized by garden, along with source population from the northern range edge (N1, N2), latitudinal range center (C1, C2), and southern range edge (S1, or S2).

### Supplementary Figures

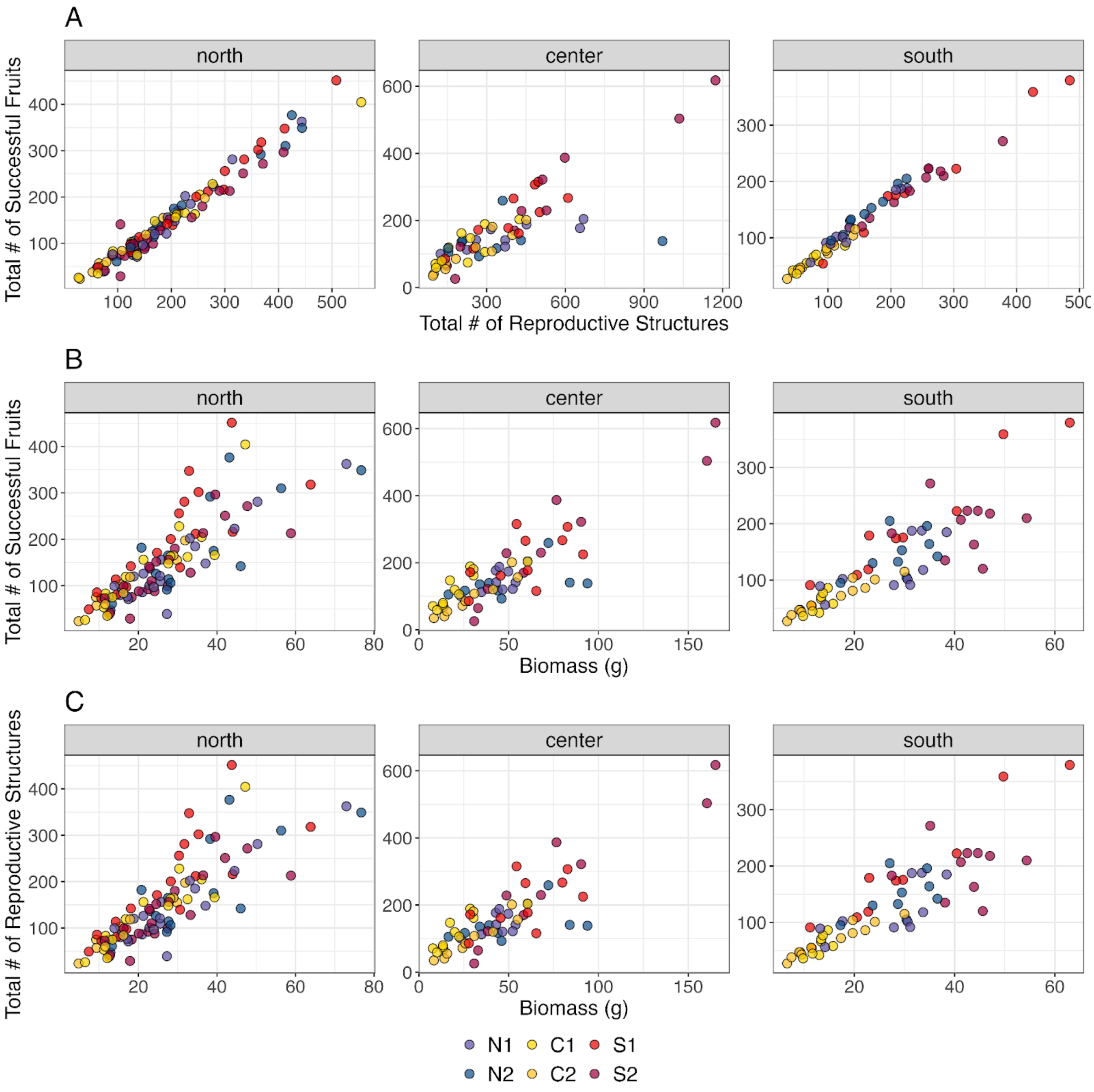


**Figure S1.** Correlations with fitness variables measured for a subset of *M. cardinalis* individuals to support the use of proxy measurements in the formal analyses of this study. Correlations between total number of reproductive structures and total number of successful fruits for plants in the northern, central, and southern common gardens (**A**). Correlations between dry biomass (g) and total successful fruits for plants in the northern, central, and southern common gardens (**B)**. Correlations between biomass and total reproductive structures for plants in the northern, central, and southern common gardens (**C**). Pearson's correlation coefficients (*r*) and their associated *P* values for each panel are found in Table S3.

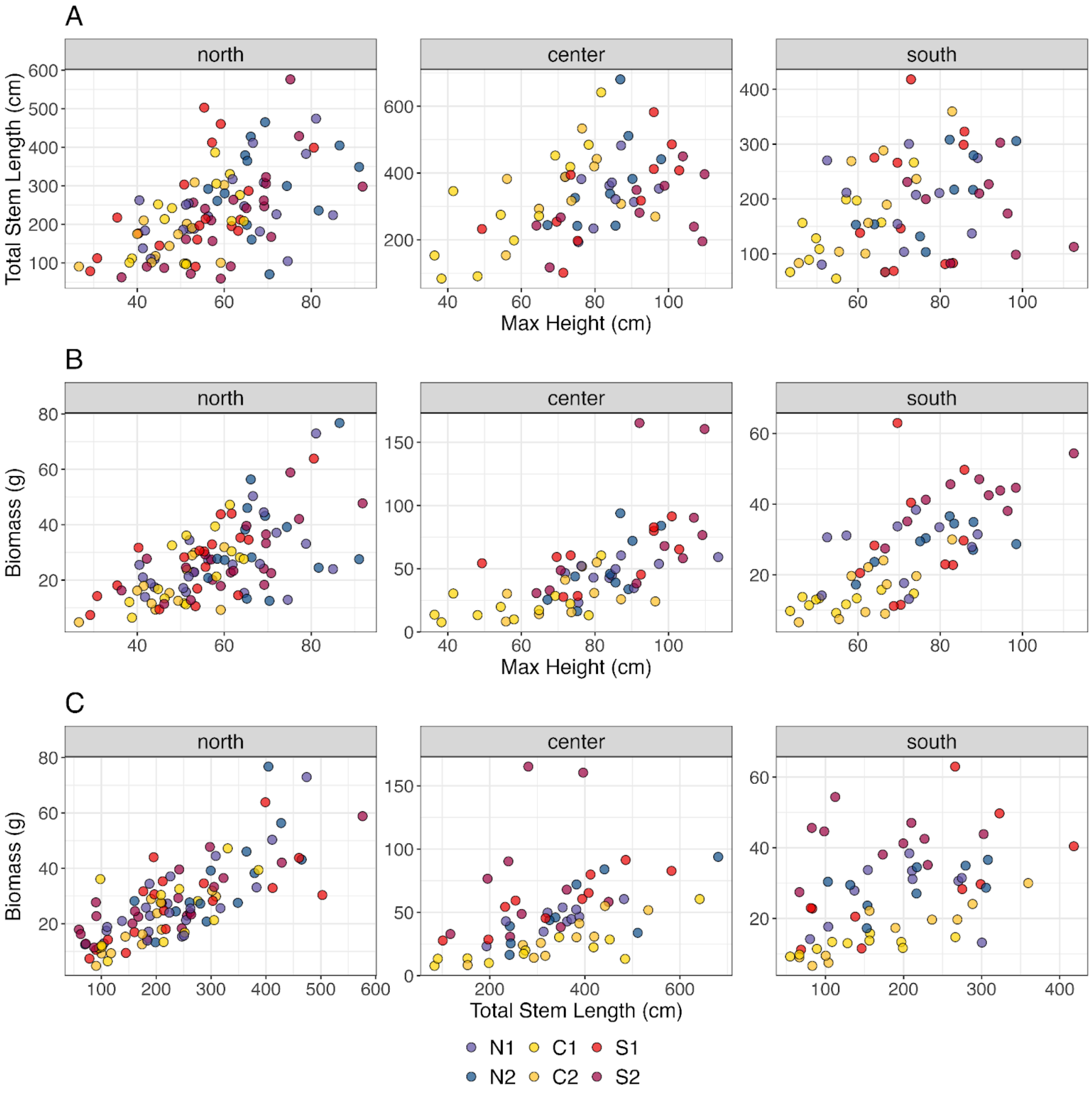


**Figure S2.** Correlations with growth variables measured for a subset of *M. cardinalis* individuals to support the use of proxy measurements in the formal analyses of this study. Correlations between length of the longest stem (cm) and total stem length (cm) for plants in the northern, central, and southern common gardens (**A**). Correlations between length of the longest stem and biomass for plants in the northern, central, and southern common gardens (**B**). Correlations between total stem length and biomass for plants in the northern, central, and southern common gardens (**C**). Pearson's correlation coefficients (*r*) and their associated *P* values for each panel are found in Table S3. N1 and N2 represent source populations from the northern range edge, C1 and C2 are from the latitudinal range center, and S1 and S2 are from the southern range edge.

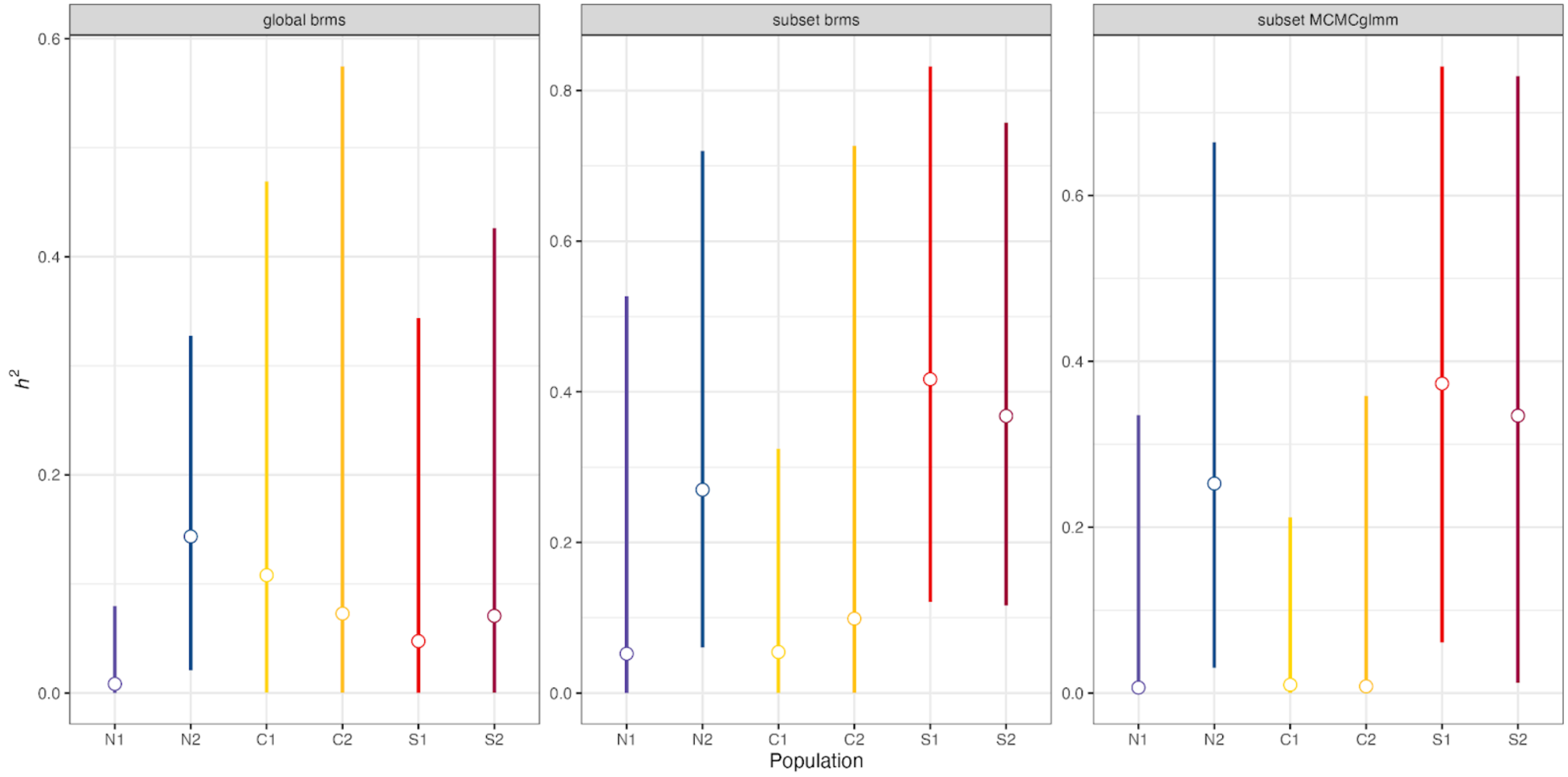


**Figure S3**. Narrow-sense heritability ($h^2$) in flowering time for the ancestral cohort of each population in in its local garden (i.e., populations N1 and N2 in northern garden, C1 and C2 in central garden, and S1 and S2 in southern garden). White circles represent median values, and lines show 95% confidence intervals. Estimates are derived from three methods, shown in different panels: 1) a global model including data from all cohorts, populations, and gardens using the brms package (left panel, as in Table S8 and Fig. 5), and models built with data from only the ancestral cohort, separately for each population in each garden using the 2) brms (center panel), and 3) MCMCglmm packages (right panel).

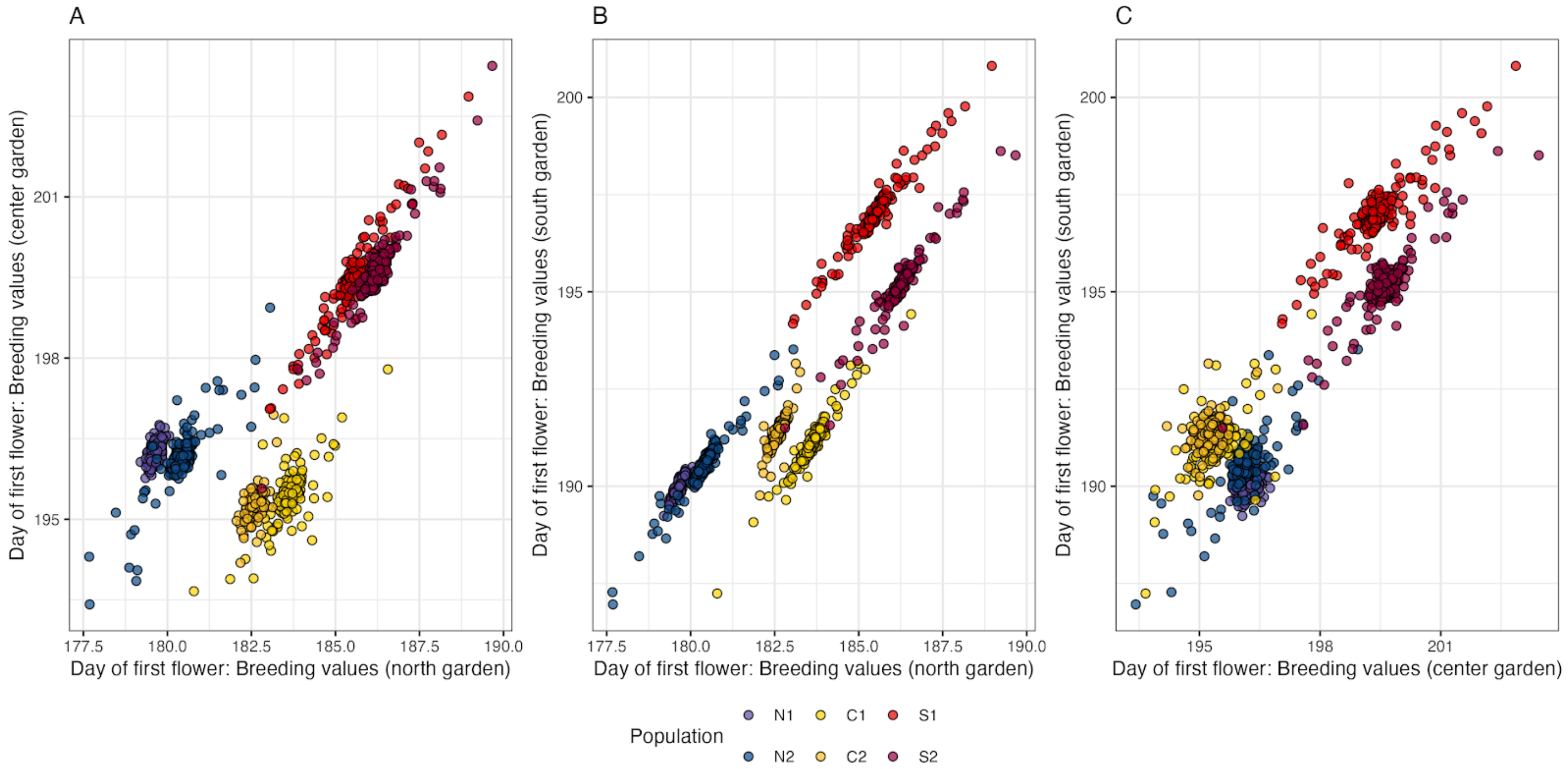


**Figure S4**. Breeding values for day of first flower for each population of *M. cardinalis* in each pair of gardens.
